# Proteasome mutations associated with CANDLE syndrome cause altered neuronal development by dysregulating polyamine synthesis

**DOI:** 10.1101/2025.08.14.670165

**Authors:** Clayton W. Winkler, Benjamin Schwarz, Katie Williams, Sara Alehashemi, Simote T. Foliaki, Joseph Snow, Lisa Joseph, Audrey Thurm, Christopher Friend, Gwendolyn Cooper, Eric Bohrnsen, Farzana Bhuyan, Nathan T. Brandes, Ruin Moaddel, Manfred Boehm, Guibin Chen, Cole D. Kimzey, Bibiana Bielekova, Joanna Kocot, Peter Kosa, Cathryn L. Haigh, Raphaela Goldbach-Mansky, Karin E. Peterson

## Abstract

Genetic mutations affecting proteasome function can result in multi-organ diseases, such as Chronic atypical neutrophilic dermatosis with lipodystrophy and elevated temperature (CANDLE) syndrome. Neurological symptoms associated with CANDLE suggest that proteasomal mutations may impact neuronal development and/or function. We generated cerebral organoids (COs) from CANDLE patient induced pluripotent stem cells (iPSCs), which exhibited impaired neuronal development when compared to COs from healthy control iPSCs. Impaired neuronal maturation in CANDLE COs was correlated with increased polyamines, which were also elevated in CANDLE patient CSF. The proteasome-regulated Ornithine decarboxylase (ODC), a rate limiting enzyme for polyamines, was elevated in CANDLE neurons. Inhibition of ODC reversed polyamine overproduction and repaired neuronal maturation in CANDLE COs, suggesting a potential therapeutic avenue for intervention. These findings demonstrate that dysfunction of the proteasome affects neuronal development through overproduction of polyamines via dysregulation of ODC and offer insight into potential therapeutic strategies for CNS-related proteasomal dysfunction.

## Introduction

Genetic mutations leading to over-activation of innate immune responses can cause autoinflammatory diseases with chronic systemic and organ specific inflammation. Neurocognitive disorders are increasingly reported in autoinflammatory diseases suggesting an important role for innate immune genes in neurodevelopment. Exploring how these genetic mutations influence the developing central nervous system (CNS) may yield valuable insights brain development and may offer potential clues for developing treatments for different neurological conditions.

*C*hronic *A*typical *N*eutrophilic *D*ermatosis with *L*ipodystrophy and *E*levated temperature (CANDLE), a recently characterized proteasome-associated autoinflammatory syndrome (PRAAS), is mostly caused by autosomal recessive founder mutations in the proteasome subunit beta type-8 gene, *PSMB8.* These mutations reduce function of the 20S proteasome core, disrupting intracellular proteolytic degradation(de Jesus et al., 2019; Montealegre Sanchez et al., 2020). Impaired proteostasis results in stress and unfolded protein responses leading in chronic type I interferon (IFN) production and inflammation, hallmark features of CANDLE/PRAAS(Brehm et al., 2015; Davidson et al., 2022). Although rare, CANDLE/PRAAS can also be caused by autosomal recessive loss of function (LOF) mutations in *PSMB9* and *PSMB4* and by digenic recessive or dominant negative forms of inheritance in subunits of the 20S core proteasome or the proteasome assembly proteins *POMP* and *PSMG2*(Brehm *et al*., 2015; de Jesus *et al*., 2019; Liu et al., 2012; Poli et al., 2018).

CANDLE/PRAAS patients typically present in infancy with systemic inflammation, including fevers and rashes due to neutrophilic panniculitis that can lead to lipoatrophy in the extremities as well as face and musculoskeletal symptoms(Sanchez et al., 2018). Patients have elevated type I IFN signatures in the blood. Suppression of IFN signaling through treatment with Janus kinase (JAK) inhibitors is effective in controlling inflammatory disease manifestations of CANDLE/PRASS(Sanchez *et al*., 2018). However, other metabolic features can persist despite treatment(Sanchez *et al*., 2018), suggesting other mechanisms also contribute to CANDLE/PRASS. Developmental and cognitive delays were reported in the early descriptions of CANDLE/PRAAS, then referred to as Nakajo Nishimura syndrome(Tanaka et al., 1993), but are not present in all cases(Torrelo, 2017). CNS manifestation commonly include basal ganglia calcifications(Garg et al., 2010; Liu *et al*., 2012; Torrelo, 2017; Tufekci et al., 2015) and in rare cases, seizures have been reported(Garg *et al*., 2010). Cognitive impairment is most pronounced in children with severe early-onset disease.

The role of the proteasome in neuronal development and function is not well understood. Proteasomes are involved in regulating axonal regeneration, calcium signaling, neuronal plasticity, and synapse number and function(Alvarez et al., 2023; Ramachandran and Margolis, 2017; Sun et al., 2019). The beta5i subunit of the proteasome encoded by *PSMB8* is expressed at appreciable protein levels in both neurons and neural stem cells, although at lower levels than in some cells such as fibroblasts(Alvarez *et al*., 2023). Recent studies suggest that several idiopathic intellectual disability cases map to mutations within the proteasome and its regulator, suggesting an important role of the Ubiquitin Proteasome System (UPS) in neurons, potentially impacting neuronal development(Cuinat et al., 2024). Thus, CANDLE-causing *PSMB8* mutations might have an underappreciated role in neuronal maturation and function and impact the CNS manifestations of CANDLE.

To study the impact of 20S proteasome dysfunction on neuronal differentiation and neurodevelopment, we generated cerebral organoids (COs) using iPSCs derived from CANDLE/PRAAS patients and healthy controls. We utilized early stage (26 day old) COs as they contain primarily developing neurons without other confounding cell types, such as glia, and offer a 3-D in vitro environment to model genetic and molecular mechanisms that cause neurodevelopmental disorders(Fleck et al., 2023; Uzquiano et al., 2022). Cellular composition and organization of COs mimic fetal human cortical development(Velasco et al., 2019) and the 3-D structure offers distinct advantages in mapping metabolic interactions and tracing cell transport and migration during neuronal development beyond existing cell culture models(Morales and Andrews, 2022). Histological and flow cytometric analyses of the COs from CANDLE (CAN) patients showed fewer neural progenitor cells and mature neurons compared to healthy control (HC), suggesting altered neurodevelopment. These cerebral organoids as well as cerebral spinal fluid (CSF) from CAN patients were analyzed for metabolic changes to identify mechanisms underlying this altered neurodevelopment.

## Results

### CANDLE syndrome patients have low Wechsler Intelligence scores

Patients diagnosed with CANDLE syndrome have reported CNS-associated pathologies including basal ganglia calcifications, intellectual disability, and in rare cases, seizures(Garg *et al*., 2010; Liu *et al*., 2012; Tanaka *et al*., 1993; Torrelo, 2017; Tufekci *et al*., 2015). We used standardized psychometric testing(Bigler, 2021) to assess neurocognition in 5 adolescence or young adult patients with clinically and genetically confirmed CANDLE syndrome. We utilized age- and ability-appropriate Wechsler Intelligence Scale instruments, to estimate intellectual abilities(Coalson et al., 2010; Watkins et al., 2022) in this cohort. Patients CAN1, CAN2, CAN5 and CAN7 were tested once, while patients CAN3 and CAN4 were tested twice. All six patients scored below low average (85 IQ) at all test dates and 4 of the 6 patients consistently scored below extremely low (≤69 IQ) (Fig. 1). These data, along with previous reports, suggest that lower than normal IQ may be a component of CANDLE/PRAAS syndrome.

**Figure 1.**
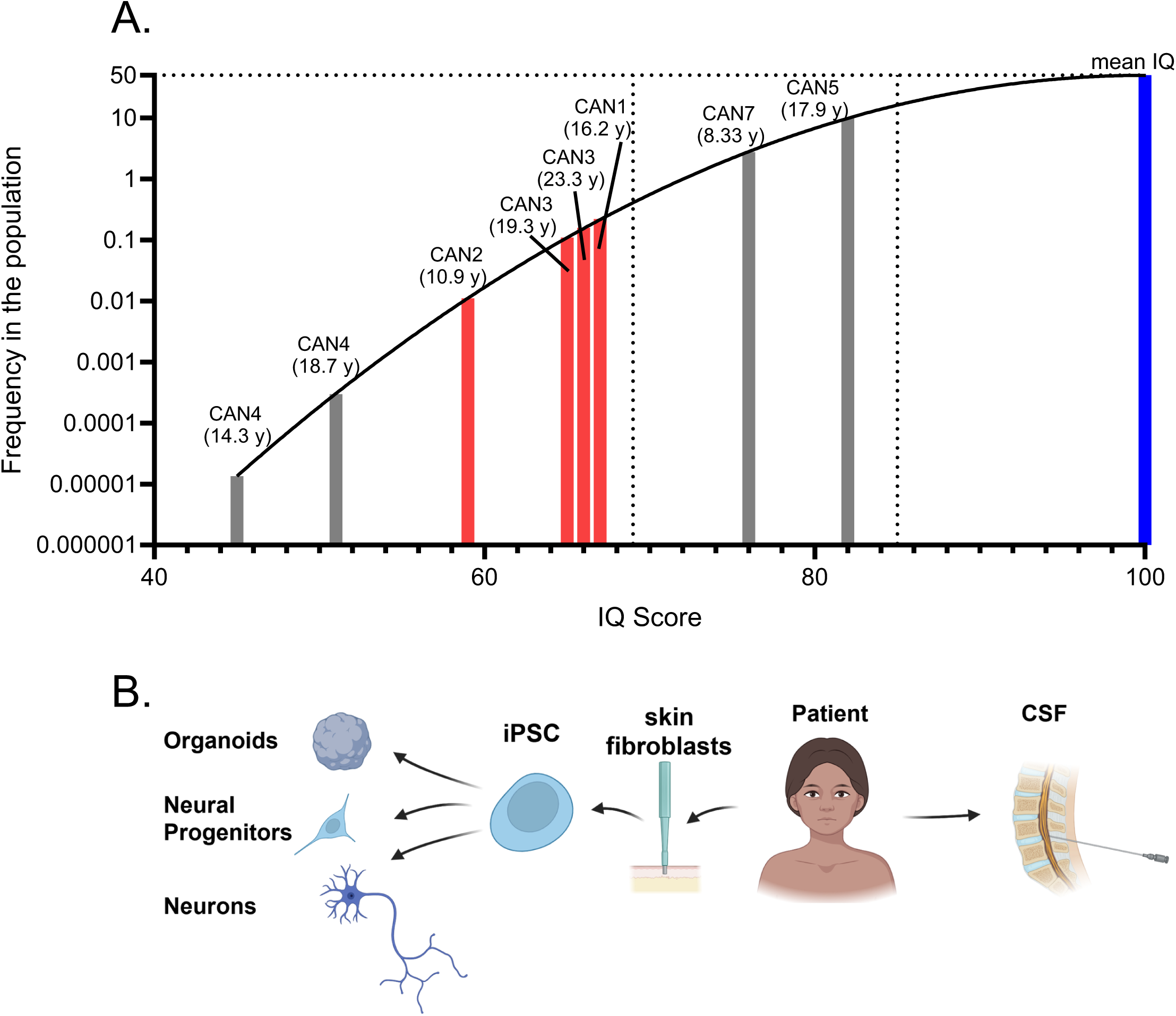
CANDLE patients have lower than average Wechsler Intelligence scores. (A) The plotted black line represents the frequency (y-axis) of an IQ score (x-axis) in a normal population with the blue bar to the right indicating an IQ of 100 (the mean standardized IQ score, SD=15). Age-appropriate Wechsler Intelligence Scale testing results of three CANDLE patients, for which iPSCs were generated were plotted as the red bars. Patients where iPSCs were not generated are plotted as gray bars. The age at each testing is given above the bar. The horizontal dotted line indicates a population frequency of 50 and the two dotted vertical lines indicate an IQ of 69 (intellectual disability) and 85 (low normal) from left to right. The results demonstrate low IQ scores for all patients, implying impaired neurodevelopment. (B) Schematic showing sample collection, processing, tissue generation and analysis from CANDLE patients. For patients CAN1, CAN2 and CAN3, skin biopsies were used to generate iPSCs from skin fibroblasts. These iPSCs were then utilized to generate cerebral organoids, neural precursor cells, and neurons. CSF was used from patients CAN3, CAN6 and CAN7 for metabolomic analysis. There was no IQ score for CAN6.

### COs generated from CANDLE patient iPSCs have altered neurodevelopment

To examine how CANDLE/PRASS syndrome impacts neurodevelopment, we generated COs from induced pluripotent stem cells (iPSCs) from three CANDLE patients (CAN1, CAN2, CAN3) and two healthy controls (HC1 and HC2) (Fig. 1B, Table S1). All CANDLE patients had *PSMB8* mutations; CAN2 and CAN3 were homozygous or compound heterozygous for pathogenic *PSMB8* variants and CAN1 had digenic variants in *PSMB8* and *PSMA3* (Table S1). Although the HC iPSC donors were both older than the CAN donors at the age of iPSC generation, previous studies on donor age indicated that age at generation did not substantially impact iPSC differentiation potential, senescence or functionality(Strässler et al., 2018).

Between 12-22 COs per each iPSC line were analyzed by immunohistochemistry at 26 days post neural induction, a time point where COs consistently contain a developmental spectrum of neuronal cells, from progenitors to mature neurons, without the confound influence of astrocytes. COs generated from HC donors showed typical cellular organization with SOX2^+^ neural progenitors forming rosette-like structures throughout the organoid (Fig. 2A) and TUJ1^+^ projections, indicative of mature neurons, (Fig. 2A-B) filling the space between the progenitor-filled structures. In contrast, COs from CANDLE patients had overall less dispersal of SOX2^+^ rosettes throughout the tissue (Fig. 2C) and less TUJ1^+^ label throughout the organoid (Fig. 2D). Quantification of SOX2 (Fig. 2E) and TUJ1 (Fig. 2F) signal as a percent of CO area was lower in CANDLE COs compared to HC COs, with the average for each CAN patient being lower than the average of either HC. Flow cytometric quantitation of SOX2^+^ and TUJ1^+^ cells on individual COs generated from CAN1 and HC1 iPSCs (Fig. S1) showed that the percentage of TUJ1^+^ cells were significantly lower in CAN COs compared to HC COs (Fig. 2G). Thus, neuronal cell maturation was altered in developing CANDLE COs relative to HCs suggesting CANDLE proteasomal mutations influence neurodevelopment.

**Figure 2.**
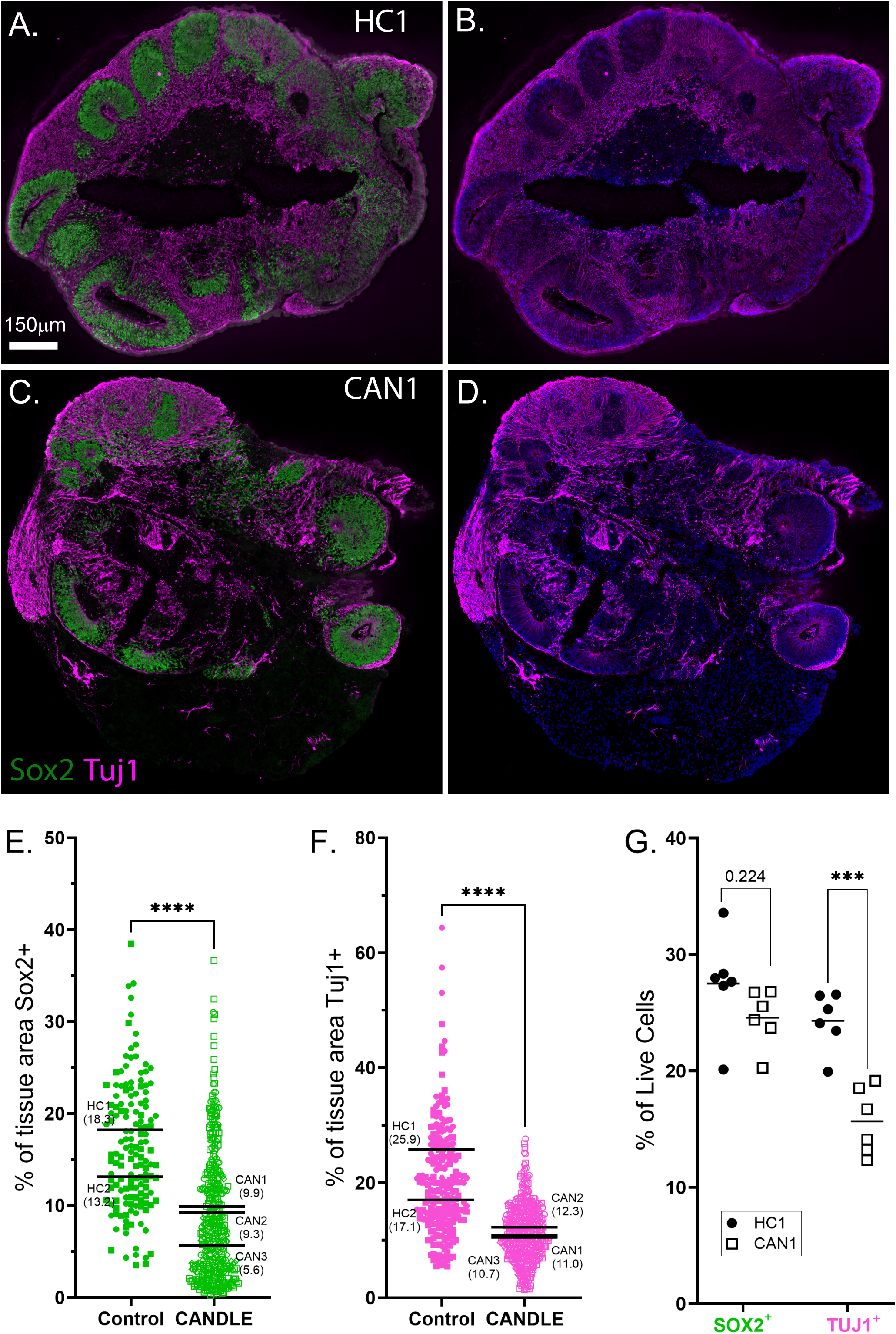
CANDLE COs have altered development. (A-F) Immunohistochemical analysis of neural progenitor (Sox2) and mature neuron (Tuj1) markers in (A, B) HC1 and (C, D) CAN1 cerebral organoids. Blue signal is cell nuclei. Quantification of immunohistochemical labeling of (E) Sox2 and (F) Tuj1 analyzed on sections from n=56 CANDLE and n=31 healthy control organoids. Data are plotted as percent of each signal within each tissue section using symbols described in Table S1 to indicate COs from generated from different iPSCs. An unpaired t test between each group was performed. **** indicates p<0.0001. **(G)** Flow cytometry was performed on n=6 organoids generated from HC1 and CAN1 with labeling for the same Sox2 and Tuj1 markers. Data is shown as the percentage of positive live cells from whole dissociated organoid. A 2way ANOVA with multiple comparisons was performed. *** indicates p<0.001. Other p values are indicated with specific values.

### CANDLE does not induce dysregulated IFN response in COs

As CANDLE mutations are associated with chronic IFN overproduction(de Jesus et al., 2020; Ebstein et al., 2019; Sanchez *et al*., 2018), we analyzed IFN and cytokine production in HC and CAN COs. Surprisingly, no difference in gene expression of *IFNB1*or interferon stimulated genes (ISGs) *ISG15, IFIT1* and *IFIT2* was observed between CAN and HC COs (Fig. 3A). We next analyzed supernatants from cultured HC1 and CAN1 COs, but no significant increase in production of IFN or other cytokines were observed (Fig. 3B). We then assessed the IFN response to a viral infection by inoculated COs from HC1 and CAN1 iPSC lines with one of two orthobunyaviruses, La Crosse virus (LACV) or Inkoo virus (INKV), which are known to infect neurons and induce IFN responses(Winkler et al., 2019). Surprisingly, CAN1 COs had lower *IFNβ* RNA following virus infection than HC1 COs (Fig. 3C). Likewise, the ISG response as determined by measuring *IFIT1* mRNA expression, was similar or lower between HC1 and CAN1 COs under mock or virus infection conditions. Thus, the differences in neuronal development in CANDLE COs was not associated with a heightened IFN response, suggesting another mechanism.

**Figure 3.**
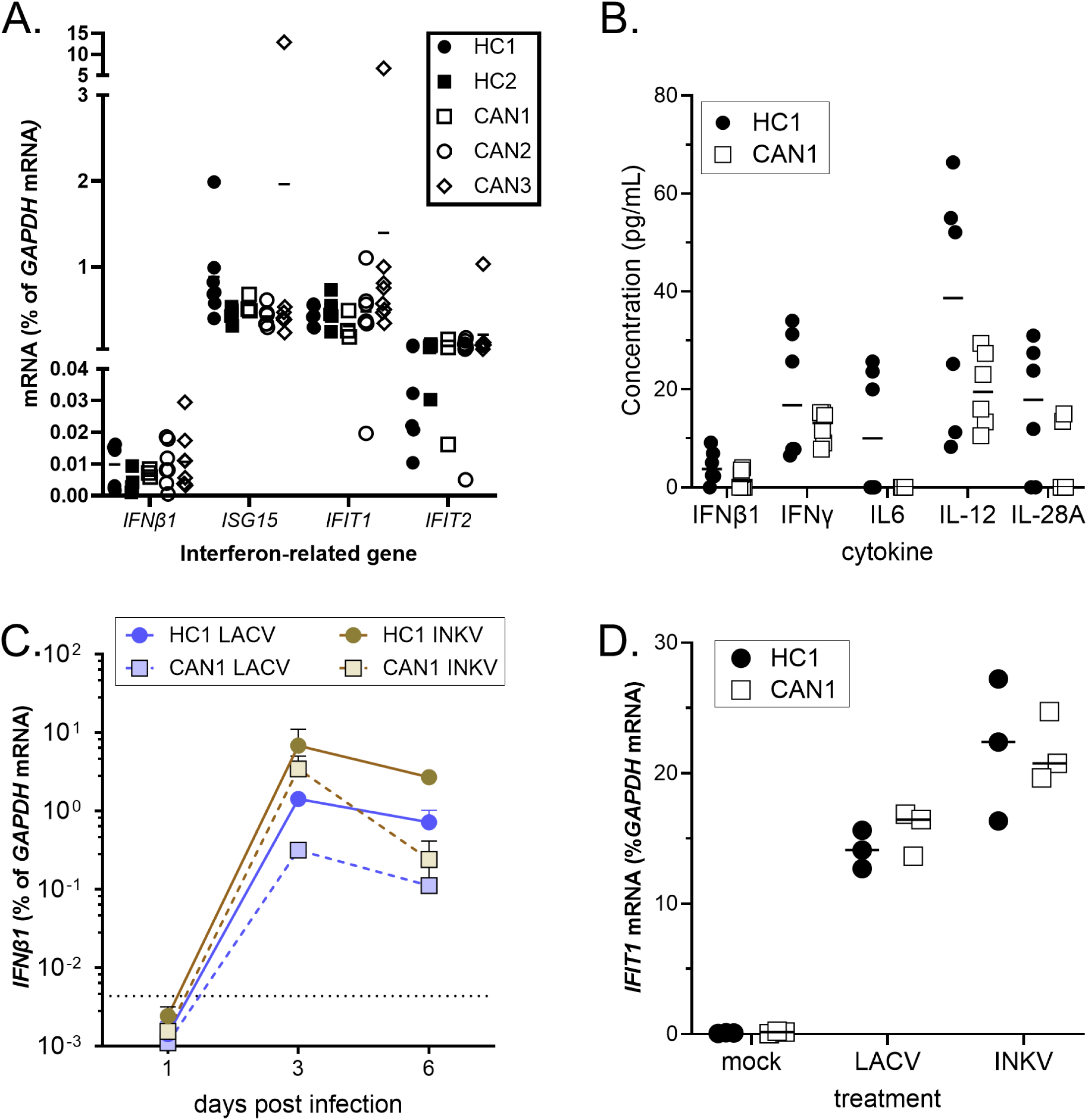
CAN COs do not have elevated cytokine or IFN responses. **(A)** cDNA reverse transcribed from mRNA isolated from COs from HC1, HC2, CAN1, CAN2 and CAN3 was analyzed by real-time PCR for the indicated genes. Data as analyzed as the ratio of *GAPDH* mRNA expression relative to the gene of interest for each organoid. A 2way ANOVA with a Sidak’s multiple comparisons test was used to compare these group. No statistical difference was observed. **(B)** Supernatants from HC1 and CAN1 COs at three weeks of age were measured for the concentration of cytokines related to the IFN response using a human inflammation 37 bio-plex multiplex immunoassay. No significant difference was observed in cytokine expression between groups. An unpaired t test between HC and CAN was used to compare groups. No statistical difference was observed. (C-D) HC1 and CAN1 COs were infected with 10^3^ PFU of La Crosse virus (LACV) or Inkoo virus (INKV). At the indicated time points, COs were collected and mRNA extracted and reverse transcribed. Individual COs were analyzed for (C) *IFNB1* and (D) *IFIT1* mRNA expression. All graphs, each symbol represents data from individual organoids or supernatants with two-to-eight samples per group per time point. An unpaired t test was used to compare HC and CAN within the mock, LACV and INKV groups. No significance was observed.

### Metabolic dysregulation associated with CANDLE in COs

Previous reports and clinical findings of significant metabolic dysregulation including protein processing, dyslipidemia and hepatic steatosis in CANDLE patients suggested that metabolic imbalances may influence cellular homeostasis and neurodevelopment(Liu *et al*., 2012; Sanchez *et al*., 2018). To examine differences in factors that may be controlling this process, we used a Biocrates mass spectrometry profiling kit to analyze metabolites from organoids as well as media from the organoids. We compared combined COs generated from HC3 to those generated from CAN1 and CAN2 iPSCs. Top differentially produced metabolites from both the cell (Fig. S2A) and supernatant (Fig. S2B) fractions included the amino acid and polyamine precursor, ornithine and the neurotransmitter gamma-aminobutyric acid (GABA), a downstream product of polyamine degradation(Sequerra et al., 2007). Both were increased in CAN1 and CAN2 COs relative to HC3 COs. The polyamine putrescine was also elevated in the cellular fraction and spermine was elevated in the supernatant of CAN COs. Thus, polyamine synthesis was elevated in CAN COs relative to HC COs as upstream, intermediate, and downstream products are all increased. These results suggested that multiple metabolic pathways were altered in CAN COs, particularly the polyamine synthesis pathway.

To expand our analysis more broadly within central metabolism, we analyzed COs from HC1, HC2, CAN1, CAN2, and CAN3 iPSCs using liquid chromatography tandem mass spectrometry (LC-MS/MS). Parallel, univariate analysis of all CO’s analyzed showed perturbation in CANDLE COs in three main metabolic pathways including amine metabolism, Acetyl-CoA associated central metabolic processes, and nucleobase metabolism (Fig. 4A, C-E). Mapping of these data onto the respective pathways showed a shift in CANDLE COs toward increased polyamine production (Fig. 4B), an increased standing pool of nucleobases (Fig. S3A), and perturbed Acetyl-CoA-generating metabolism (Fig. S3E).

**Figure 4.**
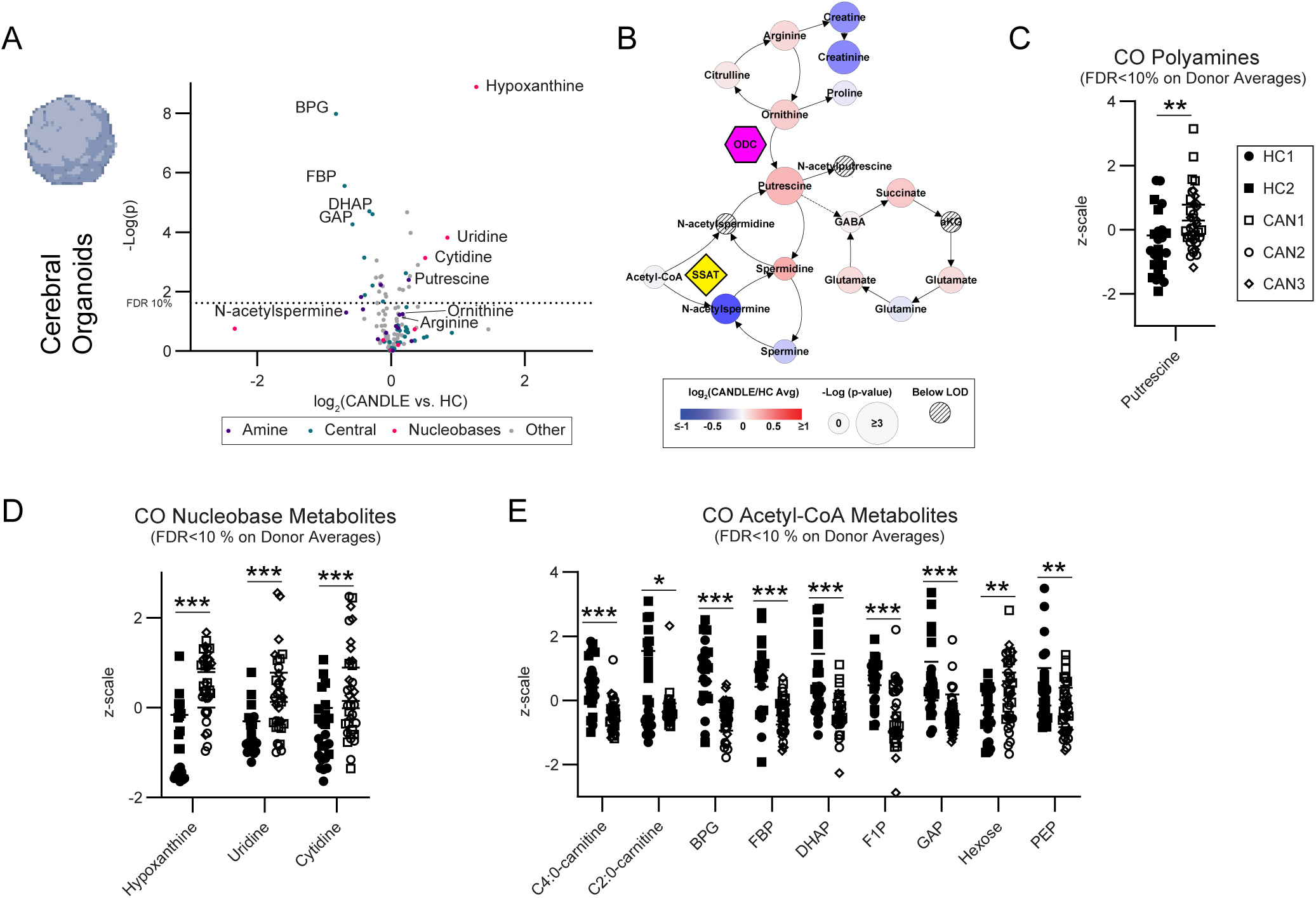
COs from CANDLE patients have altered polyamine, nucleobase and Acteyl-CoA metabolism. **(A)** A comparison of CANDLE vs. HC CO metabolites via t-test (technical n=12 per donor, biological n=3 CAN n=2 HC). Metabolites annotated as being associated with amine metabolism dark blue dots), Acetyl-CoA metabolism (light blue dots), and nucleobase metabolism (red dots) are indicated. A line representing the raw p-value cut-off for an FDR of 10 % calculated via a Benjamini-Hochberg correction is indicated. **(B)** Amine metabolism map of the same CO metabolite comparison displayed in **(A)** with the fold change of metabolites represented as color and the significance represented as node size as indicated in the legend. Hatch marked nodes indicate metabolites below the limit of detection. CO expression of key **(C)** polyamine **(D)** nucleobase and **(E)** Acetyl-CoA metabolites that passed an FDR 10 % cutoff by Benjamini- Hochberg correction. Data are shown for all assayed COs in z-scale using the variance of all samples to normalize the distribution for comparison purposes. An unpaired t test was used to compare between groups * indicates p<0.05, ** indicates p<0.01, *** indicates p<0.001.

### Metabolic dysregulation associated with CANDLE CSF

To determine if these changes in metabolite pathways were also observed in CANDLE patients, we analyzed metabolites in the CSF from three CANDLE patients, one of which (CAN3) had been used in the previous CO studies and two other CANDLE patients with available CSF as CAN6 (digenic *PSMB4, PSMB9*) and CAN7 (*PSMB8* dominant negative) (Fig 1B). These were compared to seven CSF samples from healthy controls unrelated to the CO studies as well as 10 CSF samples from Multiple Sclerosis (MS) patients to provide a contrast with an inflammatory CNS disease (Table S2). In agreement with CO data, CANDLE patient CSF showed perturbations within the same metabolic pathways, with an even more pronounced increase in polyamine and GABAnergic and nucleobase metabolites than observed in COs (Fig 5A-E, S3B, S3E). CSF from patients with active MS did not show dysregulation observed in CANDLE CSF (Fig 5C-E), suggesting these pathways alternations were due to CANDLE/PRAAS related metabolism, rather than neuroinflammation. Although two of the CANDLE CSF patients were of substantially lower age than the control HC and MS CSF (Sup. Table 2), the patient with the highest increase in amine and nucleobase changes was the oldest patient (CAN7). Additionally, although polyamine levels decrease substantially in the CNS in the first year of life, they are consistent or slightly increased in the brain from one year to middle age(Albright et al., 1983; Morrison et al., 1995). Thus, it is unlikely that age was the contributing factor for the CSF changes observed in the CANDLE patients.

**Figure 5.**
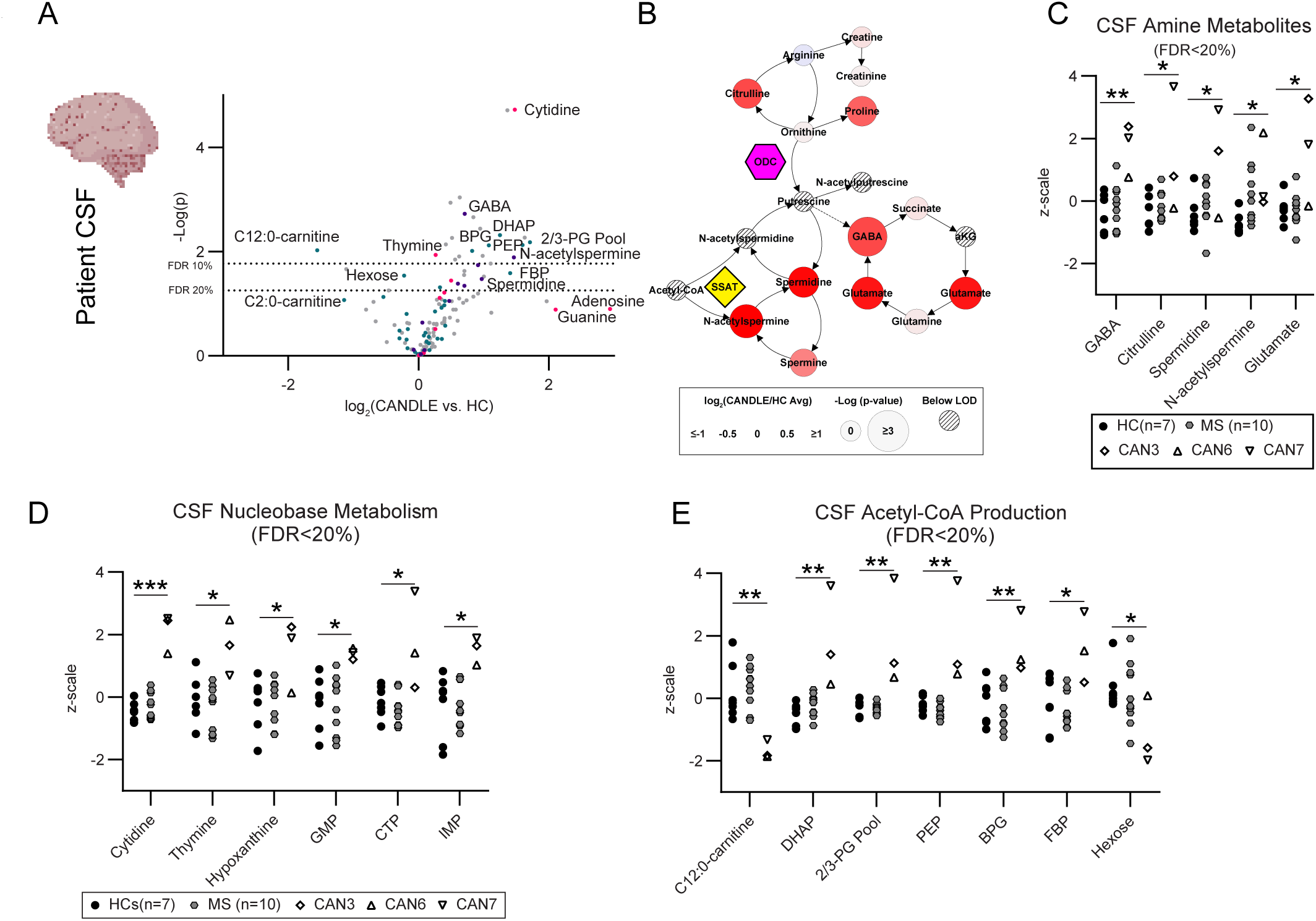
CSF from CANDLE patients have altered polyamine, nucleobase and Acteyl-CoA metabolism. **(A)** A comparison of CANDLE vs. HC CSF metabolites via t-test (biological n=3 CAN n=17 HC). Metabolites annotated as being associated with amine metabolism, Acetyl-CoA metabolism, and nucleobase metabolism are highlighted. Lines representing the raw p-value cut- off for an FDR of 10 % or 20 % calculated via a Benjamini Hochberg correction are indicated. **(B)** Amine metabolism map of the same metabolite data and comparison displayed in **(A)** with the fold change of metabolites represented as color and the significance represented as node size as indicated in the legend. Hatch marked nodes indicate metabolites below the limit of detection. Patient CSF data for **(C)** amine, **(D)** nucleobase and **(E)** Acetyl-CoA metabolites that passed an FDR 20 % cutoff are shown for each individual patient or control. Data are plotted in z-scale using the variance of all samples to normalize the distribution for comparison purposes. An unpaired t test was used to compare between groups * indicates p<0.05, ** indicates p<0.01, *** indicates p<0.001.

### Analysis of pathways dysregulated by CANDLE mutations

The pathway with the highest induction of metabolites in CANDLE COs and CSF compared to HC controls was nucleobase metabolism (Fig. 4A, 5A, Sup. Fig. 3A-B). Hypoxanthine was markedly increased in CANDLE COs and is an important indicator of nucleotide salvage pathway activity within the brain. Elevated levels of nucleobases may be due to recycling of existing nucleobases (purines and pyrimidines) due to broad dysregulation indicating CNS stress (Biasibetti-Brendler et al., 2018; Irvine et al., 2022).

The amine metabolism pathway perturbation in CANDLE COs and patients was less intense than the nucleobase effect but highly consistent between the COs and patient CSF and was found in both mass spectroscopy analyses applied to the COs (Fig. 4-5, Sup. Fig. 2). The level of polyamines did vary between CAN CSF samples, with CAN6 often within the range of the HC donors, while CAN3 and CAN7 were generally higher (Fig. 4A-F). These variations could be influenced by multiple factors, including specific CANDLE mutations (Table S1-S2). In COs, the shortest chain polyamine, putrescine, was significantly increased in CANDLE COs, while other polyamines trended towards higher levels (Fig. 4B, 4C). In CSF, both GABAnergic metabolites and longer chain polyamines and polyamine metabolites were elevated in CANDLE, including spermidine, spermine and n-acetylspermine (Fig 5B, C). Although direct interactions between nucleobase and polyamine metabolism have not been clearly established, polyamines can regulate nucleotide production and usage, suggesting that these two upregulated pathways could be connected(Zahedi et al., 2022). Together, these data suggest that in context of the CNS, CANDLE-related mutations in the 20S proteasome led to increased polyamines and nucleobases.

Polyamine synthesis is controlled by the rate-limiting enzyme in polyamine synthesis, ornithine decarboxylase (ODC) (Bewley et al., 2006). This ODC pathway of polyamine synthesis is controlled by proteasomal regulation(Bewley *et al*., 2006). Polyamine degradation, on the other hand, occurs via a separate pathway initiated by spermidine/spermine N1-acetyltransferase (SSAT), which utilizes Acetyl-CoA as a substrate for the acetylation, inactivation, and export of polyamines(Bewley *et al*., 2006). Pathways leading to the generation of Acetyl-CoA, including glycolysis and fatty acid oxidation (FAO), were the third metabolic pattern shared between CANDLE COs and CSF (Fig. S3E-F). However, CANDLE-associated effects on glycolytic and FAO metabolites differed substantially between COs and the CSF (Fig. S3E-F). This may be due in large part to the intracellular localization of glycolysis, FAO and Acetyl-CoA more generally, with the metabolites in CO samples being reflective of the intracellular metabolome, while the CSF metabolites reflect nutrients and secreted metabolic products in the extracellular space of the CNS. This contrast in the metabolite landscape of these matrices is emphasized by the expected low levels of high energy phosphorylated glycolytic compounds relative to hexose in the CSF compared to the CO (Fig S3I).

### CANDLE mutations impact polyamine metabolism differently in neural progenitor and neuronal cells

As neuronal phenotypes in COs were directly affected by CANDLE mutations, we next asked if there were CANDLE-associated metabolic differences in immature neural-progenitor cells (NPCs) and cells more differentiated towards neurons. iPSCs from HCs and patients were differentiated towards each phenotype and then analyzed for metabolic changes. As observed in CSF and COs, CAN NPCs had substantial changes in polyamines, nucleobases and Acetyl-CoA compared to HC NPCs (Fig. 6A-B, S3D, S3H). However, although CANDLE NPCs had alterations in the same pathways as observed in CANDLE COs and CSF, there were distinct differences with downregulated polyamines as well as nucleobases, rather than the upregulated responses observed in COs and CSF (Fig. 6A-B, Sup. Fig. 3D, H). For example, polyamines, including putrescine and spermine were decreased in CAN NPCs relative to HC NPCs (Fig. 6B). Interestingly, Acetyl-CoA and N-acetylspermidine were increased in CAN NPCs suggesting abundant polyamine acetylation in CAN NPCs. (Fig. 6B). Therefore, we analyzed *SSAT*, a regulator of polyamines, in one HC and one CAN NPC line. We found a trend towards higher SSAT mRNA in CAN NPCs (Fig. 6C). Thus, CAN mutations result in a metabolic signature in NPCs that is dominated by the same three pathways, but the specific patterns are distinct from those seen in the more complex organoid model or in patient CSF.

**Figure 6.**
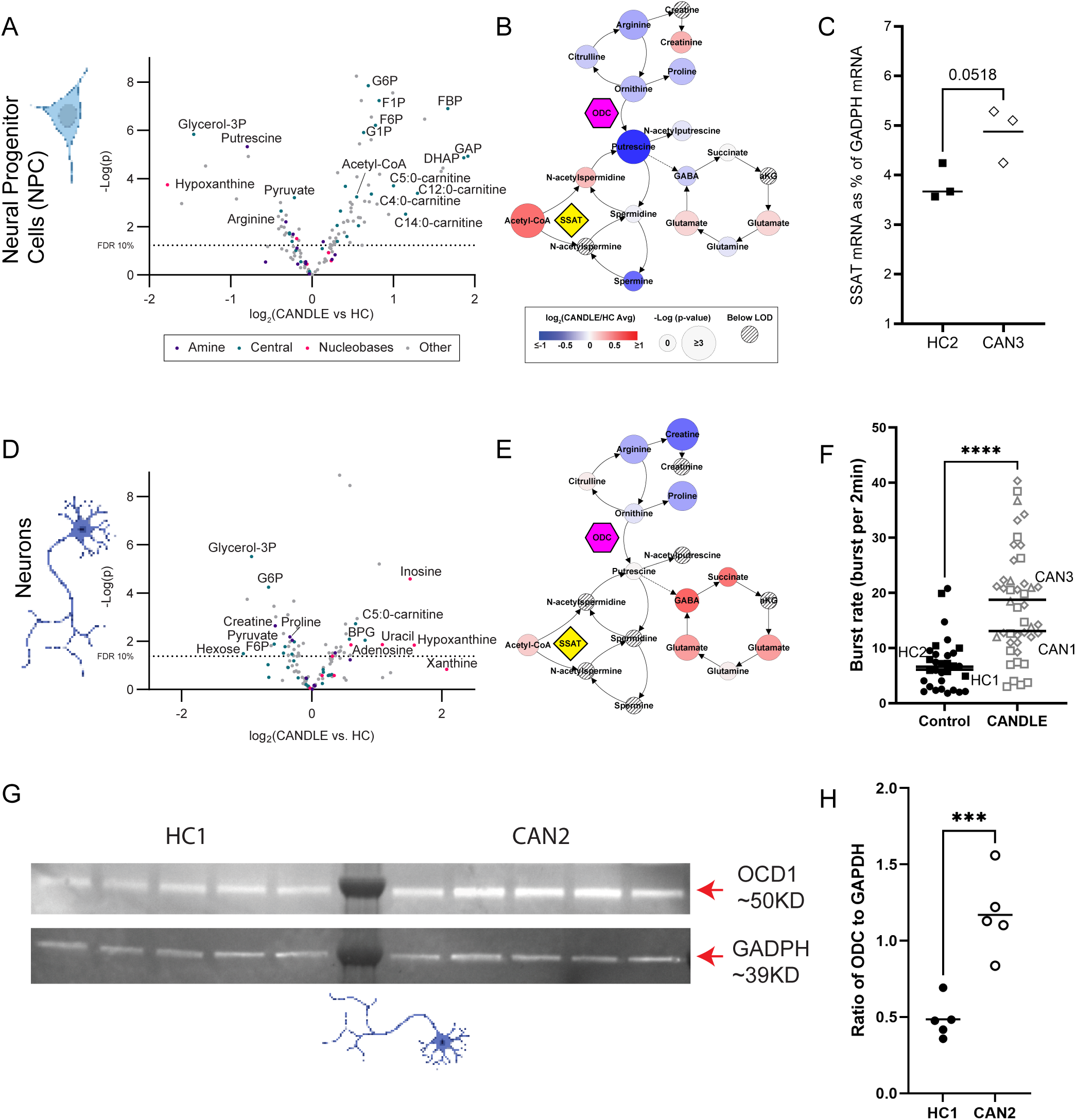
NPC and Neurons with CANDLE mutation have altered polyamine metabolism. **(A)** Comparison of CANDLE and HC NPC metabolites via t-test (technical n=6, biological n=2 CAN n=3 HC). Metabolites annotated as being associated with amine metabolism (purple dots), Acetyl-CoA metabolism (light blue dots), and nucleobase metabolism (magenta dots) are indicated. A line representing the raw p-value cut-off for a FDR or 10 % calculated via a Benjamini Hochberg correction is indicated. **(B)** Amine metabolism map of the same NPC data displayed in **(A)** with the fold change of metabolites represented as color and the significance represented as node size as indicated in the associated legend. Hatch marked nodes indicate metabolites below the limit of detection. The yellow diamond indicates Acetyl-CoA usage by SSAT for the inactivation of polyamines via acetylation. **(C)** Transcriptional expression of *SSAT* in NPCs derived from HC1 and CAN3 (n=3 replicate wells for each). Data are plotted as *SSAT* expression as a percent of *GAPDH* housekeeping gene. Data were compared by a two-tailed unpaired t-test. P value is indicated. **(D)** Comparison of CANDLE and HC Neuron metabolites via t-test (technical n=6, biological n=3 CAN n=3 HC). Metabolites annotated as being associated with amine metabolism (purple dots), Acetyl-CoA metabolism (light blue dots), and nucleobase metabolism (magenta dots) are indicated. **(E)** Amine metabolism map of the same neuron metabolite data displayed in **(D**) with the fold change of metabolites represented as color and the significance represented as node size. The yellow diamond indicates Acetyl-CoA usage by SSAT for the inactivation of polyamines via acetylation. **(F)** Burst rate defined as the number of bursts (at least three spikes detected within 100ms) over a 2min period in cultured neurons from n=2 HC and n=2 CAN (15-37 technical replicate wells recorded for each patient). The average readout of active channels per well represented a replicate. Data were compared between cell lines by a two-tail unpaired t-test. **** indicates p<0.0001. **(G)** Western Blot analysis for OCD1 expression by neurons generated from HC1 and CAN2. n=5 wells from a 12 well plate with 5x10^5 cells. The monomeric ODC is ∼50kD in molecular weight in the top blot. The same blot was reprobed for GAPDH expression at ∼39KD (bottom blot) **(H)** Ratiometric densitometry analysis of ODC expression relative to GAPDH control.

Analysis of metabolic changes in more mature differentiated neurons also showed metabolic signature dominated by changes in polyamine, nucleobase and Acetyl-CoA pathways with CAN mutations (Fig 6D-E, Sup. Fig. 3C, G). As observed in CAN CSF, CAN neurons showed increased GABAergic metabolites and glutamate metabolites, and low to undetectable putrescine (Fig. 6D-E). Hypoxanthine and pyrimidines, including cytidine, uracil, and uridine, were also increased in CAN neurons compared to HCs, in alignment with both CO and CSF samples (Sup. Fig. 3E). Acetyl-CoA directed metabolites were depleted or absent in glycolysis and FAO pathways in CANDLE neurons, distinct from COs, CSF and NPCs (Sup. Fig. 3E). Overall, the analysis of CANDLE NPCs and mature neurons showed metabolic signatures that were dominated by the same pathways as CSF and COs, indicating a consensus in affected metabolic processes. Differences in the metabolic responses between the two cell types are aligned with the expected increase in metabolic restriction with the degree of neuron differentiation (Zheng et al., 2016). The high level of neurotransmitters, including GABA and glutamate in CAN neurons (Fig. 6E) and CSF (Fig. 5B) lead us to examine if CANDLE mutations altered neuronal activity. Although GABA is normally considered an inhibitory neurotransmitter, excessive GABA can increase neuronal firing by modulating neuronal excitation to oscillate faster (Ben-Ari, 2002). Indeed, CAN neurons were more electrically active compared to HCs (Fig. 6F), suggesting CAN neurons were more active than HC neurons.

As ODC is a rate-limiting enzyme that catalyzes polyamine synthesis, we next analyzed if CAN variants had altered ODC levels. We compared expression of ODC in neurons generated from HC1 and CAN2 iPSCs. ODC was expressed in both sets of neurons, however, the intensity of the 50kD protein band was significantly higher in CAN2 neurons than HC1 neurons (Fig. 6G-H, S4). Thus, increased ODC levels was associated with CANDLE proteasome mutations and could drive the higher levels of polyamines observed in CANDLE COs and CSF.

### Inhibition of ODC in CANDLE COs reduces polyamine levels and promotes mature neuronal transcripts

To determine if blocking ODC activity would reverse the metabolic and neuronal changes associated with CANDLE, we treated HC and CANDLE COs with difluoromethylornithine (DFMO), a specific small-molecule inhibitor of ODC. DFMO containing media or media alone was added to COs at 6 days of age and replaced daily for another 20 days. Dose analysis showed that DFMO was well tolerated by COs groups and did not alter organoid growth (Fig. S5A-B). A concentration of 100μM was chosen for further studies as being the highest concentration that did not impact cellular growth (Fig. S5).

Next, HC and CAN COs were treated with DFMO for 20 days and compared to vehicle-treated controls (Fig. 7). Metabolic analysis showed a partial reversal of CANDLE-associated metabolic patterns in amine, Acetyl-CoA, and nucleobase metabolism (Fig. 7, S6 compared to S2). As expected, putrescine was downregulated relative to vehicle controls for CANDLE COs (Fig. 7B). Metabolites within the nucleobase scavenging pathway were also largely decreased in DFMO-treated CANDLE COs suggesting that polyamine metabolism was the causal driver of these nucleobase patterns in CANDLE models and patient samples (Fig. 7, S6B). Of note, the key metabolites affected by DFMO in CANDLE COs were unaffected or directionally distinct in DFMO-treated HC COs including Acetyl-CoA, FAO metabolites, hypoxanthine, and putrescine (Fig 7B-C). Thus, DFMO-treatment of CANDLE COs resulted in a substantial reversal of the metabolic pathologies associated with the CANDLE mutations in COs, but not HCs. This lack of a response in HC COs suggested that other polyamine regulatory mechanisms may buffer against DMFO’s affects in HC but not CANDLE COs.

**Figure 7.**
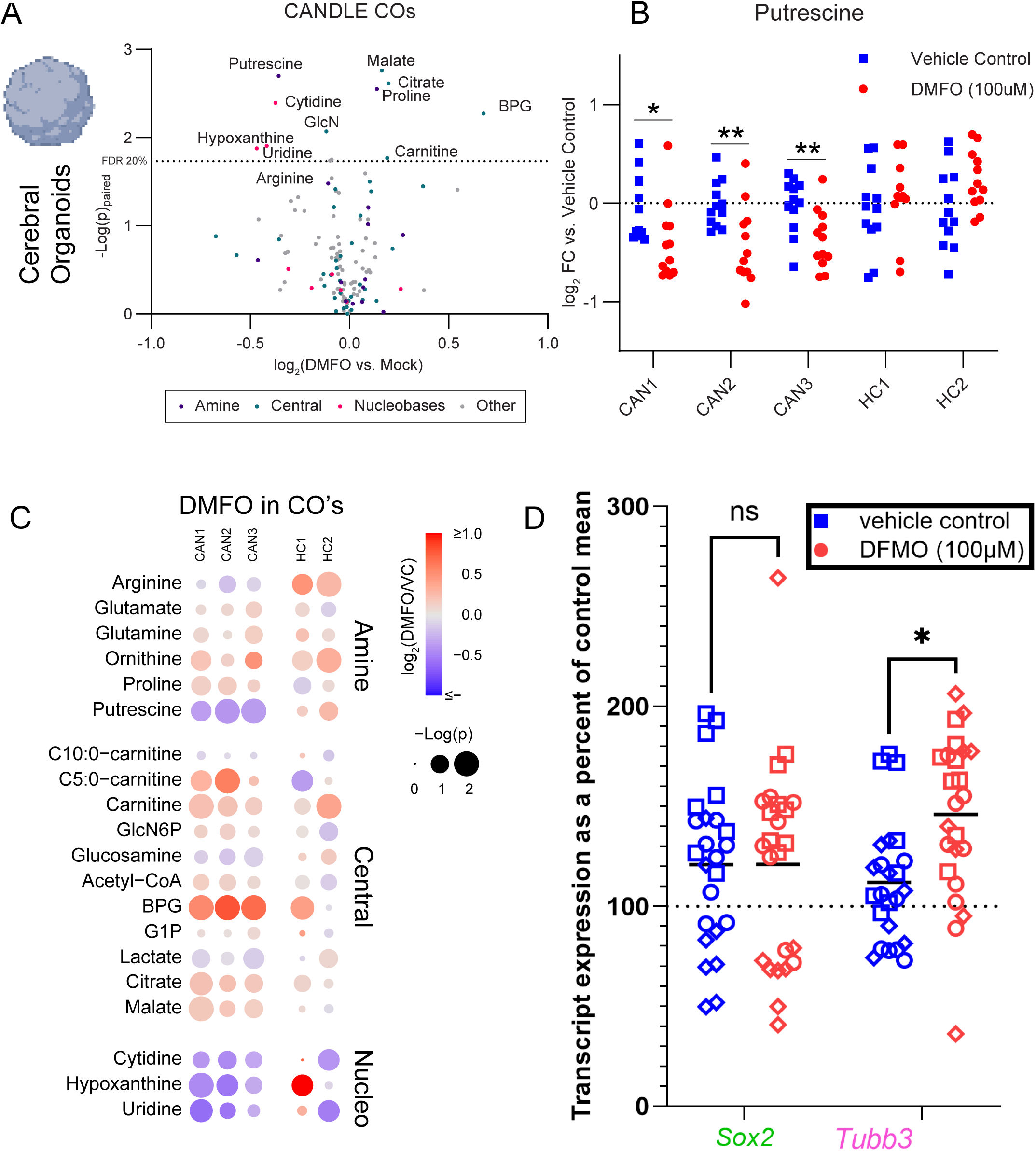
Impact of DFMO on metabolism and neuronal development in CANDLE COs. **(A)** Comparison of 100 µM DFMO-treated CAN COs vs. vehicle-treated CAN CO metabolites via paired t-test by on the average metabolite levels (CO technical n=12, biological n=3 CAN). Metabolites annotated as being associated with amine metabolism (purple dots), Acetyl-CoA metabolism (light blue dots), and nucleobase metabolism (magenta dots) are indicated. **(B)** Change in putrescine signal in COs relative to the average of the vehicle control by donor. An unpaired t test was used to compare groups. * indicates p<0.05, ** indicates p<0.01). **(C)** Metabolites that changed in response to DFMO treatment in CANDLE COs via a paired t-test with p<0.1 organized by pathway. Node color reflects the fold change due to DFMO and the size reflects the significance of that change by patient as indicated in the associated legend. **(D)** Transcriptional expression of *Sox2* a and *Tubb3* (Tuj1) with and without DFMO treatment from n=16 healthy control and N=24 CANDLE COs. Data are plotted as transcription expression as a percent of *Gapdh* housekeeping gene relative to the mean of healthy control percent of *Gapdh*. Data were analyzed by a 2way ANOVA with a Sidak’s multiple comparisons test. * indicates p<0.05.

To directly determine if DFMO reversed the phenotype of altered neurodevelopment in CANDLE COs, we analyzed CANDLE COs with and without DFMO treatment for neuronal marker expression by real-time PCR. *SOX2* mRNA, a marker of more immature stem cells was not altered by DFMO treatment (Fig 7D). However, *TUBB3* mRNA, encoding TUJ1 protein, was significantly increased across all CANDLE COs treated with DFMO relative to media control-treated COs (Fig. 7D). Thus, inhibition of polyamine synthesis promoted the expression of transcripts associated with mature neuron development in CANDLE COs.

## Discussion

In the current studies, we found that the 20S proteasome regulates maturation of neurons. This regulation was not through the expected type I IFN responses, associated with CANDLE/PRASS mutations, but rather through increased nucleobase and polyamine synthesis caused by accumulation of ODC, a protein regulated heavily by the proteasome. Targeting ODC, by treating with DFMO, reversed or ameliorated both the altered metabolism and the abnormal neuronal development. Thus, this study provides a tissue-specific mechanism by which mutations in the proteasome can affect the CNS and contribute to CANDLE/PRASS syndrome. Whether proteasomal regulation of ODC impacts polyamine regulation in other tissues remains to be determined.

Our study reveals a strong correlation with polyamine levels and neuronal maturation, consistent with other recent studies linking polyamine dysregulation and neurodevelopmental disease(Bachmann et al., 2024; Rajasekaran et al., 2021). For example, Bachmann-Bupp syndrome, associated with neurodevelopmental delays and autism, is caused by de novo gain-of-function mutations in ODC1 leading to constitutive activation of ODC and polyamine overexpression(Bachmann *et al*., 2024). Symptoms associated with Bachmann-Bupp syndrome can be partially corrected by administration of DFMO early in life(Rajasekaran *et al*., 2021). Monogenic LOF mutations in spermine synthase (SMS) causes an X-linked intellectual disability disorder, Snyder-Robinson syndrome, due to increased polyamine synthesis suggesting a possible dose effect of polyamine production on neurodevelopment(Akinyele et al., 2024). Polyamines are thought to influence cellular development by directly binding nucleic acids, remodeling chromatin, interacting with intracellular RNA, and/or regulating cell cycle progression(Igarashi and Kashiwagi, 2010; Miller-Fleming et al., 2015; Ohnuma and Harris, 2003). Given the broad pathogenic implications of dysregulated polyamine pathways on neurodevelopment, treatment with ODC inhibitors such as DFMO provides an attractive option for downregulating polyamine synthesis(Rajasekaran *et al*., 2021). In our studies, DFMO not only normalized polyamine synthesis but also corrected nucleobase and Acetyl-CoA pathways, supporting polyamine overproduction as being upstream of these cell and microenvironment-specific rescue pathways (Fig. 7). Thus, DFMO treatment may resolve polyamine dysregulation without impacting these associated pathways in healthy cells.

The CNS manifestations of CANDLE 20S proteasome mutations differ from those observed in patients with disease-causing variants affecting the 19S regulatory protein complex (19S RP) of the 26S proteasome. Those patients have more severe neurodevelopment and social behavior abnormalities within the autism spectrum(Ebstein *et al*., 2019) with little or no systemic inflammation. Within the 26S proteasome, the 19S RP unfolds and directs ubiquitinated proteins for degradation by the 20S core of the 26S proteasome, involving both 19S RP and 20S subunits for turnover of ubiquitylated proteins(Liu and Jacobson, 2013). Disruption of either subunit can impact neurons. For example, disruption of 19S RP or 20S subunits induce cellular inclusions detrimental to neurons(Droggiti et al., 2011; Rideout and Stefanis, 2002). Both subunits may also impact ODC turnover in the cell. ODC degradation can be ubiquitin-independent and executed by the 20S subunit alone(Asher et al., 2005). However, ODC activity and degradation are also controlled by the protein antizyme (AZ), whose expression is dependent on the 19S subunit for recognition of ubiquitination and subsequent degradation(Kahana et al., 2005). Thus, in 19S RP mutations, ODC levels and subsequent polyamine production may be increased similarly to our observations in 20S mutations through impaired control of AZ expression. Side-by side comparison of 19S RP and 20S mutations on neuronal development may ultimately be needed to determine the impact of the two subunits on ODC, polyamine synthesis and neuronal development.

Although the three CANDLE-associated metabolic patterns (amine, nucleobase, and Acetyl-CoA metabolism) were all observed in CSF, COs, neurons and NPCs, there were variations in the individual metabolites affected between these models, with certain metabolites highly impacted in some systems and below detection limits in others (Figs. 4-6). A primary reason for these variations may simply be the contrasting sample matrices. For example, the CSF is extracellular, whereas COs, neurons and NPCs reflect the intracellular metabolome. As most energized metabolism occurs intracellularly(Fuller and Kim, 2021), precise diagnosis of metabolic pathology from an extracellular fluid such as CSF is difficult. Indeed, there was a lower relative abundance of phosphorylated intermediates in the extracellular CSF compared to the primarily intracellular COs (Fig. 5). Another contributor to the differences in CANDLE associated metabolic patterns between both the CSF, COs, neurons and the NPCs is the metabolic restriction of the CNS at the blood brain barrier, which limits metabolic inputs including both glucose and fatty acids from the vasculature (Panov et al., 2014). Additionally, neurons are highly metabolically restricted and will avoid both direct metabolism of fatty acids and aerobic glycolysis due, in part, to the potential for oxidative damage, while both FAO and glycolysis are known to be active and essential in NPCs(Knobloch et al., 2017; Zheng *et al*., 2016). We found cell-specific metabolic patterns in NPC and neuron models that suggest the magnitude of the CANDLE-associated response in Acetyl-CoA directed pathways, which provide substrate for polyamine inactivation via acetylation, is much greater in NPCs compared to neurons (Fig. 6, S3). Thus, the buffering capacity to cope with excess polyamines via degradation is anticipated to be greater in NPC because of the metabolic diversity available to these cells. In contrast, in metabolically restricted neurons, polyamines may be shunted toward alternative processes such as the GABAnergic cycle. These restrictions in Acetyl-CoA metabolism in the CNS, as well as the direct functional consequences of polyamines entering the GABAnergic cycle, likely contribute to the pleiotropic effects of polyamine dysregulation on CNS disease pathology(Kilb and Kirischuk, 2022; Kovács et al., 2022; Schönfeld and Reiser, 2013). The specific location, severity, and cell types affected may provide context in other CNS disorders that have been associated with polyamine dysregulation including Alzheimer’s, schizophrenia, depression, and epilepsy(Baroli et al., 2020). The full scope of the effect of polyamine dysregulation on other neural cells and how these mechanisms all contribute to CANDLE-associated pathologies remains to be elucidated.

Treatment of CANDLE COs with the ODC inhibitor DFMO increased transcript levels for TUJ1 (*TUBB3*), but not *SOX2* mRNA (Fig. 6D). The proteasome influences protein translation in the brain through regulated degradation of translation initiation and inhibition protein factors, such as Fragile X mental retardation protein (FMRP)(Baugh and Pilipenko, 2004). Under normal conditions, FMRP binds to specific mRNA species, possibly including *SOX2*(Brighi et al., 2021; Luo et al., 2010) and silences their translation(Richter et al., 2015). Thus, the impaired proteasomal degradation in CANDLE may lead to decreased protein translation through increased mRNA silencing. This could explain why we did not observe changes in *SOX2* mRNA levels following DFMO treatment (Fig. 7D) despite finding decreased protein in untreated COs (Fig. 2E). Alternatively, the more immature SOX2^+^ progenitor cells may be less impacted by DFMO due to less stringent metabolic restrictions compared to the mature TUJ1^+^ cells, where the effect of controlling ODC may be more pronounced. Although not possible to do with the complexity of the organoid, a direct metabolic comparison between these two cell populations isolated from the organoid would parse this difference in response to DFMO.

Consistent with reports that brain nucleotide synthesis relies heavily on nucleobase salvage rather than de novo synthesis(Dobolyi et al., 2011; Sekine et al., 2024), increased purine and pyrimidine salvage inputs including hypoxanthine, uridine, cytidine, and uracil were seen in all CANDLE models tested, except in NPCs. Hypoxanthine-stimulated ROS production has been proposed as a mechanistic link of this metabolite to brain stress and injury and stress(Frenguelli and Dale, 2020; Mink and Johnston, 2007). However,xanthine oxidoreductase (XOR), which generates ROS from hypoxanthine, is minimally expressed in brain tissue negating the possibility for hypoxanthine driven ROS production(Frenguelli and Dale, 2020; Mink and Johnston, 2007). Instead, increases in hypoxanthine, and other nucleobase salvage inputs such as uridine, may be correlated with CNS stress while functioning to support increased nucleotide triphosphate demand in situations of high neural activity such as seizures(Frenguelli and Dale, 2020). This is evident in patients’ CSF and iPSC-derived neurons, where increased adenosine accompanies rises in hypoxanthine, uridine, uracil, and cytidine suggesting the use of hypoxanthine for ATP generation via adenosine. Regardless of the mechanistic impact of nucleobases on CNS stress, CANDLE-driven effects on nucleobase metabolism were downstream of ODC dysregulation, with DFMO reducing nucleobase levels in CANDLE COs. While a specific linkage between polyamines and nucleobases is possible, the association may be indirect with excess polyamines causing stress, leading to increased nucleotide recycling to prevent nucleotide depletion and cell death(Foliaki et al., 2020; Sekine *et al*., 2024).

In conclusion, our studies with cerebral organoids, CSF, and neuronal cell culture indicate that mutations in the 20S proteasome impair neurodevelopment through a lack of ODC degradation leading to increased polyamines, increased nucleobase metabolism, and dysregulated Acetyl-CoA pathways. These pathways may provide further insight into the potential mechanisms of CNS pathology associated with CANDLE and provide potential targets for therapeutic intervention with drugs such as DFMO. As well, comparison of these results with other autoinflammatory diseases, including other proteasome-related diseases, may provide a better understanding of the mechanisms that drive brain development and how dysregulation of these responses lead to brain pathology.

## Methods

### Experimental model and study participant details

All research investigations were done as part of natural history protocol 17-I-0016/NCT02974595 and as such were approved by the NIAID Institutional Review Boards. The patients were referred to the NIH between 2007 and 2013. All patients and their parents provided consent or assent for study enrollment and the study procedures including IQ testing, skin biopsies, iPSC generation and lumbar punctures. The study was reported according to CARE guidelines and conducted in compliance with the Declaration of Helsinki principles. Patient intelligence quotient (IQ) testing was compared to age-matched norms reflected in standardized scores for each test. Patient age at testing, sex/gender and disease-associated mutations are provided in Table S1. Induced pluripotent stem cells (iPSCs) were generated from fibroblasts isolated from patient skin biopsies (Yu et al., 2022). The HC1 iPSC line was obtained commercially from ATCC (https://www.atcc.org/products/acs-1023), while the generation of HC2 iPSCs has been previously described as being generated from a skin punch biopsy and reprogrammed using ReproRNA™-OKSGM (Foliaki *et al*., 2020).

### IQ testing

Cognitive function was assessed with the use of the following age-appropriate standardized tests: Wechsler Adult Intelligence Scale 4^th^ edition or Wechsler Intelligence Scale for Children 5^th^ Edition and whole IQ scores are reported in Figure 1.

### Virus Culture

As previously described, La Crosse virus (LACV) stocks derived from a 1978 human case were generated by infecting confluent Vero cells (ATCC) in a T75 flask (Corning) with an LACV multiplicity of infection of 0.01, and harvesting the supernatant from the flask after 5 days of culture (Winkler et al., 2024). Plaque forming units (PFU) of virus stocks were determined by plating 10-fold serial dilutions onto Vero cells in confluent 24-well plates (Corning). Cells were cultured in 2% FBS and penicillin/streptomycin (Gibco) containing DMEM (Gibco) media. Plates were incubated for 1h to allow virus attachment and then each well was overlayed with 500mL of 1.5% carboxymethyl cellulose (Sigma) in MEM (Gibco). Plates were incubated five days, then fixed for 1-2 hours with 10% formaldehyde (Sigma) to a final concentration of >4% formaldehyde. Plates were then rinsed with in house de-ionized water (diH2O) and stained for 10min with 0.35% crystal violet made up in 20% ethanol (Sigma) and in house diH2O. Plates were rinsed, air dried, and the numbers of plaques counted. Titers were calculated as the number of plaques per 1mL of supernatant applied.

### iPSC culture

iPSCs were maintained in reduced growth factor Matrigel (Corning)-coated 25cm^2^ culture flasks in mTeSR Plus media (StemCell Technologies) with frequent culture splitting (∼every 5 days) to maintain ≤70% confluency to minimize cellular differentiation. Splitting involved detaching cells with enzyme-free cGMP ReLeSR (StemCell Technologies) for ≤3min to leave differentiated cells attached to the flask and transferring replicating iPSCs to a new flask. iPSCs were fed daily with mTeSR Plus to maintain high levels of fibroblast growth factor in the culture to inhibit differentiation.

### Neuroprogenitor cell (NPC) and mature neuron generation

NPC cultures were generated using the STEMdiff^TM^ SMADi Neural Induction kit (StemCell Technologies) and the monolayer culture protocol with a plating concentration of 2×10^5^ cells/cm^2^. After passage 3 (18-21 days post induction), cells were either transferred into STEMdiff^TM^ Neural Progenitor Medium (StemCell Technologies) for ongoing culture of NPCs or differentiated into neurons. For neuronal differentiations, cells were plated at a density of 1×10^5^ cells/cm^2^ at passage 3 of NPC differentiation in STEMdiff^TM^ SMADi Neural Induction Medium. After 24 hours, the media was removed and replaced with STEMdiff^TM^ Forebrain Neuron Differentiation Medium (StemCell Technologies), and this media was changed daily for 6 days. After the 6 days of neuronal differentiation, cells were replated at a density of 5×10^4^ cells/cm^2^ in STEMdiff^TM^ Forebrain Neuron Maturation Medium (StemCell Technologies) and cultured for a minimum of 14 days to allow maturation. Medium was changed every three days or as required.

### Cerebral organoid generation, culture and drug administration

Cerebral organoids (Cos) were generated from CANDLE (CAN) patient and healthy control (HC) iPSCs using the CO differentiation kit (StemCell Technologies) which uses the media formulations and protocol described by Lancaster and Knoblich (Lancaster and Knoblich, 2014). Specifically, 5000 iPSCs were plated into each well of a 96-well plate with 50μl of Organoid Formation Media containing 5μM dihydrochloride and cultured for 5 days with an additional 50μl of media (without dihydrochloride) being added on days 2 and 4 to form embryoid bodies. On day 5, embryoid bodies were transferred to a new 96 well plate containing 150μL of neural induction media per well for a further 2 days. The start of of neural induction (day 5) represents the “birthday” of each organoid. On day 7 (2-days old), neural-induced embryoid bodies were embedded into 20μL of Matrigel (Corning) which was then allowed to polymerize at 37°C for 30min and subsequently transferred to 2-3mL of Expansion media per 10-15 embryoid bodies in a 6 well plate. On day 10 (5-days old), Matrigel-embedded organoids to be used for metabolic or immunohistochemical analysis were transferred to a 125mL Erlenmeyer flask with 25mL of Maturation media for an additional 30 days (25-days old for metabolic analysis) or 31 days (26-days old for immunohistochemistry) respectively. Media was replaced every 4-5 days. On day 30, organoids that underwent metabolic analysis were transferred to individual wells in a 24 well plate in 1mL of Maturation media where they were cultured for 24 hrs prior to tissue and supernatant collection and processing on day 31 (26 days old). Organoids used for immunohistochemistry were fixed in 10% neutral buffered formalin on day 31 (26 days old) for further processing.

Certain HC and CAN COs were treated with varying concentrations of difluoromethylornithine (DFMO) to determine drug toxicity (Sup Fig. 4.) and to modify polyamine synthesis (Fig. 7). DFMO was dissolved directly in maturation media at 100mM and indicated dilutions were made using the same media.

For drug treatments, individual COs were transferred to individual wells within a 24 well plate on day 11 (6-days old) and 1mL of either maturation media control or maturation media containing DFMO was added to each well. Media was refreshed daily for each organoid until day 31 (26-days old) when tissue and supernatant was collected for either metabolic or transcriptomic analysis.

### Organoid infection

COs were infected as previously described (Winkler *et al*., 2019). Specifically, HC and CAN COs were transferred to individual wells in a 24 well plate as described above and Maturation media containing either 10^3^ PFU of LACV, INKV or mock Vero cell supernatant per 1mL of media were added at 21 days post- neural induction for 2h with shaking at 85rpm in the incubator. 1mL of fresh Maturation media was then added to each well for 30min to wash off virus and subsequently replaced with an additional 1mL of fresh media for future culture. COs were incubated for 6 days post infection (dpi), with culture supernatants being collected at 2, 4 and 6 dpi. At 6dpi, uninfected mock control, INKV and LACV COs had whole RNA collected as described below in the RNA isolation section. Uninfected mock controls were used as a baseline measurement of IFN responses in LACV and INKV infected COs .

### Organoid viability assay

Organoids treated with normal maturation media or different concentrations of DFMO were analyzed for cellular reductive capacity using a PrestoBlue™ assay (ThermoFisher). This is a resazurin-based assay that serves as an indicator of cellular viability. Following maturation media removal, individual organoids were incubated for 30min with 250μL of 1XPrestoBlue diluted into maturation media from a 10X stock. 50μL of each sample was read in triplicate in a flat-bottom 96 well plate (Corning) using fluorescence excitation at 560nm and emission at 590nm on an BioTek Synergy 4 (Agilent) plate reader. 1X PrestoBlue media incubated without an organoid was used as a media blank control. Baseline measurements for each organoid were taken 24hr prior to drug treatment and then readings were taken every 3 days following DFMO administration out to 21 days post treatment for toxicity studies (Sup Fig. 3A&B) or 15 days for metabolic and transcriptomic analysis (Sup Fig. 3C&D). Data are reported as percent 590nm fluorescence relative to a pretreatment baseline.

### Bioplex Assay

Analysis of cytokine/chemokine concentrations was performed on undiluted supernatant from uninfected mock controls, INKV and LACV infected CO cultures using the Bio-Plex Human Inflammation Panel 37-plex (BioRad) per the manufacturer’s protocol. All data was collected on a Luminex 200 Instrument System (Luminex) and concentrations in range of detection were determined by Standard Curve analysis in Microsoft Excel.

### Metabolomics Sample Extraction

For all liquid chromatography tandem mass spectrometry (LC-MS/MS) methods, LCMS grade solvents were used. All CO and biofluid samples were directly immersed in 0.4 mL of ice-cold methanol. NPCs and neurons in culture were first washed with 1 mL of 0.9 % NaCl, prior to the addition of 0.4 mL ice-cold methanol. Cells were then scraped to suspend all debris and transferred to microtube for further processing. To each sample 0.4 mL of water and 0.4 mL of chloroform were added. Samples were shaken for 30 minutes under refrigeration and centrifuged at 16k xg for 20 min. 400 µL of the top (aqueous) layer was collected and diluted as necessary in 50 % methanol in water for LC-MS/MS analysis.

### Metabolomics (Liquid Chromatography Tandem Mass Spectrometry)

Tributylamine was purchased from Millipore Sigma. LCMS grade water, methanol, isopropanol, formic acid, and acetic acid were purchased through Fisher Scientific. Aqueous metabolites were analyzed using two analytical methods with opposing ionization polarities (Groveman et al., 2023; McCloskey et al., 2015). Both methodologies utilized a LD40 XR UHPLC (Shimadzu Co.) system for separation and a 6500+ QTrap mass spectrometer (AB Sciex Pte. Ltd.) for detection. In negative mode methodology, samples were separated with a Waters™ Atlantis T3 column (100Å, 3 µm, 3 mm X 100 mm) and eluted using a binary gradient from 5 mM tributylamine, 5 mM acetic acid in 2% isopropanol, 5% methanol, 93% water (v/v) to 100% isopropanol over 5 minutes. Two distinct multiple reaction monitoring (MRM) ion pairs in negative mode were used for each metabolite where available. In positive mode methodology, samples were separated using a Phenomenex Kinetex F5 column (100 Å, 2.6 µm, 100 x 2.1 mm) and eluted with a gradient from 0.1 % formic acid in water to 0.1 % formic acid in acetonitrile over 5 minutes.

### Biocrates Targeted Metabolomics Assay

A targeted metabolomic assessment was carried out following the manufacturer’s protocol using Biocrates’ MxP^®^ Quant 500 kit (www.biocrates.com; Biocrates, Innsbruck, Austria), in organoids (20 µl). For 6 organoids, 300 µl of 85% Ethanol and 15% PBS was homogenized in 2 ml tubes from the Precellys Lysing Kit, soft tissue homogenizing CK14 (Bertin, Rockville, MD). The tubes were then homogenized for 2 cycles of 20 s at 6500 rpm with 30 s interval between the homogenization steps using the Precellys 24 between 0-4°C. Afterwards the tubes were centrifuged with a benchtop centrifuge and the solution was transferred to a separate 1.5 ml Eppendorf tube, where they were centrifuged for 10 mins at 17,000-x g in the cold room and the samples were stored at -80°C until analysis.

The Biocrates MxP 500 was run using a Nexera HPLC system (Shimadzu) coupled to a 6500+ QTRAP^®^ mass spectrometer (AB Sciex) with an electrospray ionization source as previously described (13). Data were quantified using WebIDQ™ software and normalized to internal quality controls. Any metabolite that had greater than 30% above limit of detection (LOD) was excluded, while those with less than 30% LOD had their values imputed as 1/5 the lowest concentration.

### Immunoblotting and analysis

iPSC-derived neurons from HC1 and CAN2 were plated into a 12 well plate at 5x10^5^ cells per well. Lysates were generated from individual wells by addition of cold 250μL of RIPA buffer (ThermoFisher) with 1X Halt™ Protease Inhibitor (ThermoFisher) for 5min followed by gentle trituration of the solution. Lysates were collected in tubes on ice and then spun at 15,000g for 15min to remove debris. A Peirce™ BCA analysis was performed on all samples and 5μg of total protein was loaded per well into a Criterion XT 10% Bis/Tris acrylamide gel (BioRad). Proteins were electrophoretically separated by SDS-PAGE at 200V for 50min then transferred to a Immun-Blot® PVDF (BioRad) membrane via wet transfer in 1X Tris-Glycine at 100V for 1hr. The membrane was rinsed twice in tris-buffered saline (TBS) for 10min and blocked in blocking buffer (TBS with 5% milk (BioRad) and 1% TritonX100 (Sigma-Aldrich)) for 1hr at room temperature (RT) with 80rpm shaking. Primary antibodies against ODC (Abcam, 1μg/mL) or GAPDH (Novus, 5ng/mL) diluted in blocking buffer were applied overnight at 4°C with 80rpm shaking. The membrane was rinsed 3x5 min in TBS-T and alkaline phosphatase-conjugated secondary (Goat α Rabbit or Goat α mouse, 1:1000) was applied for 2hr at RT with 80 rpm shaking. The blot was them rinsed 3x15min in TBS-T and AttoPhos® Substrate (Promega) was applied for 5min. Blot was visualized using the chemiluminescent setting on an iBright Imaging System (ThermoFischer)

### Immunohistochemistry

At 26 days old, whole COs were place in 10% neutral buffered formalin for 3-7days. COs were prepared for cryosectioning by rinsing 3x1hr in 1XPBS and incubating in 30% sucrose in 1XPBS at 4°C overnight. Organoids were then embedded into OTC compound (Sakura) contained within a 15x15mm cryomold (EMS) and frozen on block dry ice. 10μm sections were generated on a Leica CM3050S cryostat (Leica Biosystems) and maintained at -20°C until used. Sections to be labeled were warmed to RT, residual dry OTC was removed, and sections were outlined with a Dako hydrophobic barrier pen (Agilent) for on-slide immunolabeling. Sections were washed 3x5min with 1X PBS and blocked for 30min at RT with 5% normal donkey serum (Millipore Sigma). Rabbit anti-Sox2 (Abcam, 1:1000) or mouse anti-Tuj1 (B3T, Millipore Sigma, 1:500) primary antibody was applied in blocking buffer overnight at 4°C. Sections were washed 3x5min in 1XPBS and AlexaFluor secondary antibodies donkey anti-Rabbit 488 or donkey anti-mouse 594 (Thermo Fisher, 1:1000) in 1X PBS were applied for 2hrs at RT. Sections were washed once with 1X PBS and Hoechst 33342 (Thermo Fisher, 0.1μg/mL) dissolved in PBS was applied for 15min. Sections were washed 3x5min in 1X PBS and slides were coverslipped with Prolong™ Diamond antifade mountant (Thermo Fisher) and #1.5 cover glass (Ted Pella). Sections were images in their entirety using a Leica Versa Slide Scanner (Leica) with a HC Plan APO 20x objective (NA 0.75).

### RNA isolation and quantitative reverse transcription (qRT) PCR from COs

At 26 days old, whole COs were collected in a new nuclease-free tube and all media was removed. COs were dissociated by adding 0.5mL of Corning™ Cell Recovery Solution (Corning) to each tube and incubating for 1hr on ice with intermittent (∼every 15min) gentle trituration. Dissociated tissues were washed with 1X PBS 3 times by pelleting the sample at 500g and replacing with fresh supernatant. After a final spin, 600μL of ZR RNA buffer (Zymo) was added to the sample and gentle trituration was used to dissolve the cell pellet. Whole RNA was isolated and according to the manufacturer’s Quick-RNA Midiprep kit protocol with the exception that the DNAse digestion was performed at 37C for 30min using Ambion™ DNAse I (RNase-free) reagent. RNA concentrations were determined spectrophotometrically using a Nanodrop One (ThermoFisher) and RNA (100ng) was reverse transcribed to cDNA using an iScript reverse transcription kit (Bio-Rad) per the manufacturer’s instructions. Reactions were performed on a MasterCycler nexus GX2 (Eppendorf) using a 5 min priming at 25°C, 30 min RT reaction at 42°C, a 5 min RT inactivation at 85°C and a 20 min stabilization at 20°C. cDNA was diluted 5-fold in nuclease-free water prior to us real-time PCR analysis.

Primers to detect human *Gapdh*, *Sox2*, *Tubb3*, *SSAT, IFNβ1, IFIT1, IFIT2,* and *ISG15* were designed using the NCBI primer design tool (NCBI) and confirmed to be specific for the target gene by blasting against the NBCI database. Primer sequences are shown in the Key Resources Table. All primers were tested on cDNA derived from normal organoids and confirmed to amplify a single target by dissociative melt curve analysis. SYBR Green dye with ROX (Bio-Rad) was used for all real-time PCR analysis per the manufacture’s protocol. Samples were run in a 384 plate in triplicate on a QuantStudio Pro7 Real-time PCR system. No reverse transcriptase and water only controls were used. Data from each sample was calculated as the percent difference in Cq value with the housekeeping gene *Gapdh* (ΔCq=(Cq *Gaph*)- (Cq gene of interest)).

### Flow cytometry

At 26 days old, whole HC1 and CAN1 COs were collected and dissociated with Cell Recovery Solution as described above. A single cell suspension was generated by passing the dissociation solution through a 70μm filter (Corning). In a 5mL tube, fixable viability dye eFluro™ 780 (Invitrogen, 1:1000) in 1X PBS with 0.04% BSA was applied for 30min on ice, in the dark to all cells except an unlabeled control aliquot. Cells were then distributed into a 96 well plate for further processing, including wells designated for unlabeled, single channel, and fluorescence minus one controls. Cells were washed 2x by pelleting at 500g for 3min, removing the solution by flicking the plate and resuspending cells in 1X PBS with BSA. Cells were then fixed, permeabilized and labeled with primary conjugate antibodies against Sox2 (Biolegend, 1:50) and Tuj1 (Abcam, 1:125) with the eBioscience™ Foxp3/Transcription Factor Staining Buffer Set as indicated by the manufacturer. Briefly, cells were incubated in fixation/permeabilization working solution overnight at 4°C, spun and flicked and then permeabilization buffer with Fc receptor blocking Truestain FcX (Biolegend, 1:1000) was added for 15min on ice in the dark. Cells were spun and flicked and then permeabilization buffer containing primary antibodies was added to designated wells for 30min on ice in the dark. Cells were washed 1x with permeabilization buffer and then 2x with1X PBS by spinning and flicking. After the final wash, cells were resuspended in 200μl of 1X PBS and analyzed by a BD FACSymphony (BD Biosciences). Gating layout is shown in Supplemental Fig. 1.

### Electrophysiological recording of iPSC-derived neurons

Neuronal population synchronous firing was recorded as described previously using the multi-well multielectrode arrays (Multiwell-MEA head stage with integrated amplifier from Multichannel Systems) with 12 gold electrodes (700 μm spacing; 100 μm of electrode diameter) per well (Foliaki *et al*., 2020). Neurons were monitored for activity starting at 7 days post-plating, and the baseline activity was recorded at 2-3 weeks post-plating when network bursts became largely synchronous. The medium (BrainPhys from StemCell Technologies) was changed at least 8 hours before recording. The local field potentials or raw signals were recorded using Multiwell-System software (V.2.0.11.0 from Multichannel Systems) at a sampling frequency of 20 KHz for 2 minutes, filtered with a second-order Butterworth high-pass filter with a 300Hz cutoff and a fourth-order Butterworth low-pass filter with a 3500Hz cutoff.

### Analysis of immunohistochemistry performed on COs

Immunohistochemistry using Sox2 and Tuj1 was performed on sections from HC1 (n=15), HC2 (n=19), CAN1 (n=22), HC2 (n=19), and HC3 (n=12) individual organoids as indicated above. A total of 229 sections from HC COs and 426 sections from CAN COs were imaged using the same capture settings for each label on the Leica Aperio slide scanner as described above. Both label channels were captured in the same imaging session into a single RGB image. Using ImageJ analysis software (NIH), the Sox2, Tuj1 and Hoechst 33342 channels were split using the “Split Channels” function into individual images, converted to 8-bit gray scale images and saved. Using the Hoechst channel image, a freehand selected ROI was generated for each imaged organoid and saved to be applied to the associated single channel images to define the tissue area. If the tissue was excessively damaged or contained large regions devoid of Hoescht label, that tissue was removed from the analysis. Each 8-bit single channel image was made binary under the process tab to generate a mask of positive label. The ROI generated for the Hoechst label was then overlayed onto the Sox2 or Tuj1 masked image to define the tissue area. The Analyze Particles function was then used with the following settings to define the percent of CO tissue area positive for each label: Size (pixel^2) =1-Infinity, Circularity=0.00-1.00, Show=Masks, with the “Display results” and “Summarize” tabs selected.

### Analysis of LCMS-derived metabolic data

All signals were visually inspected for presence and integrated using SciexOS 3.1 (AB Sciex Pte. Ltd.). Metabolites with multiple MRMs were quantified with the higher signal to noise MRM. Filtered datasets of the negative mode aqueous metabolites were total sum normalized after initial filtering. The positive mode aqueous metabolomics dataset was scaled and combined with the negative mode aqueous metabolite dataset using signals for metabolites common to both methods including Arginine, Tyrosine and Threonine. Goodness of the scaling was assessed by Principle Components Analysis. A Benjamini-Hochberg method for correction for multiple comparisons was imposed where indicated in text or Fig. captions. Technical powering was used across donors for CO-related metabolomics statistics.

### Analysis of iPSC-derived neuron electrophysiology

A Multiwell-Analyzer (V 2.0.6.0 from Multichannel Systems) was used to extract electrophysiology parameters, including spikes or neuronal population firings, bursts, and burst spike rate. Spikes were detected as those above 4.0 standard deviations of the mean noise. The neuronal population burst was defined as at least three spikes detected within 100ms, and the network burst was defined as at least two channels displaying overlapping bursts. Active channels were determined as those with at least 15 spikes (with amplitudes higher than 10 μV) per minute, and active wells were those with at least three active channels. The average readout of active channels per well represented a replicate. Data were compared between cell lines by a two-tail unpaired t-test. Outliers identified by the ROUT method using Prism (v10.4.1) were excluded from the statistical comparison.

## Key Resource Table

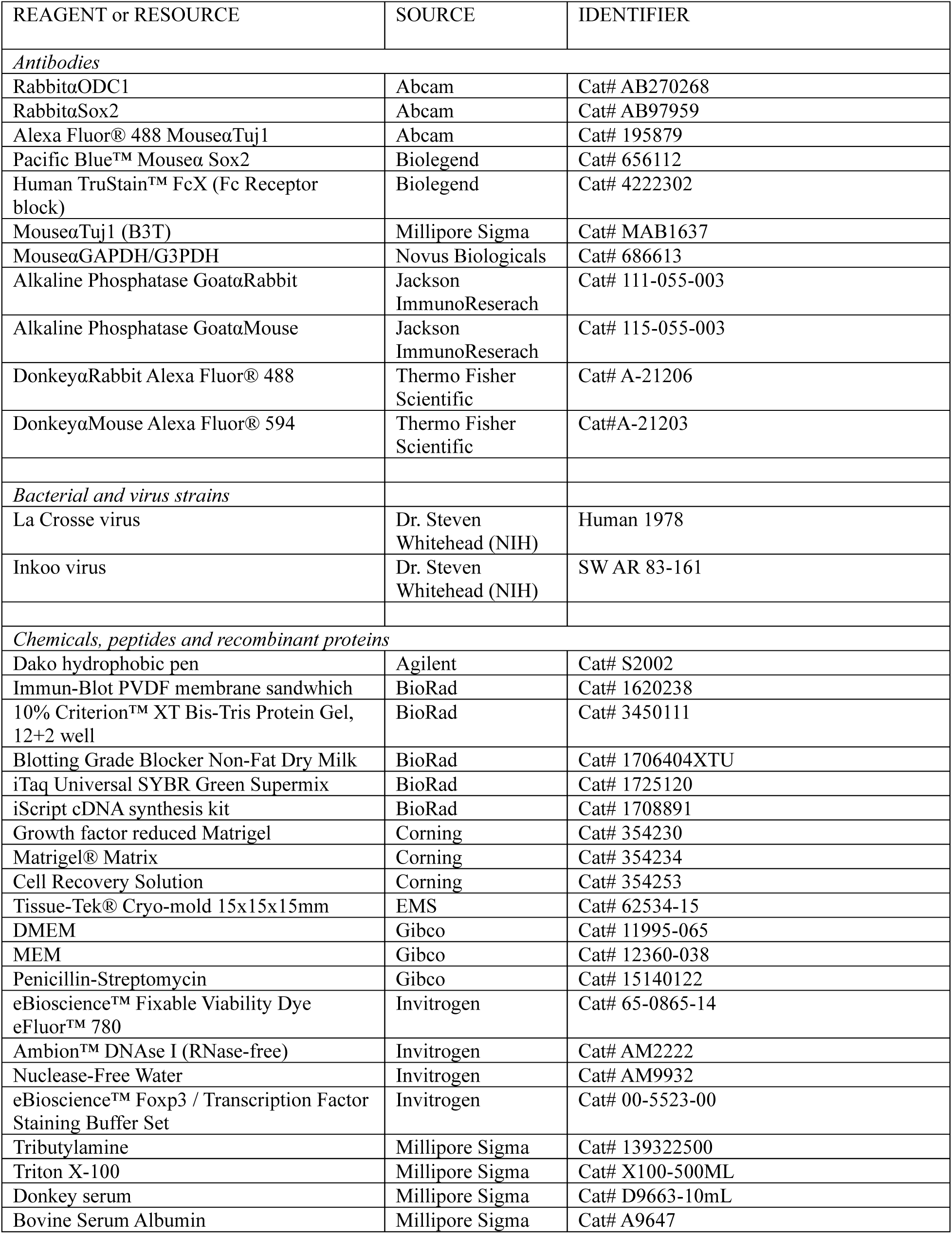
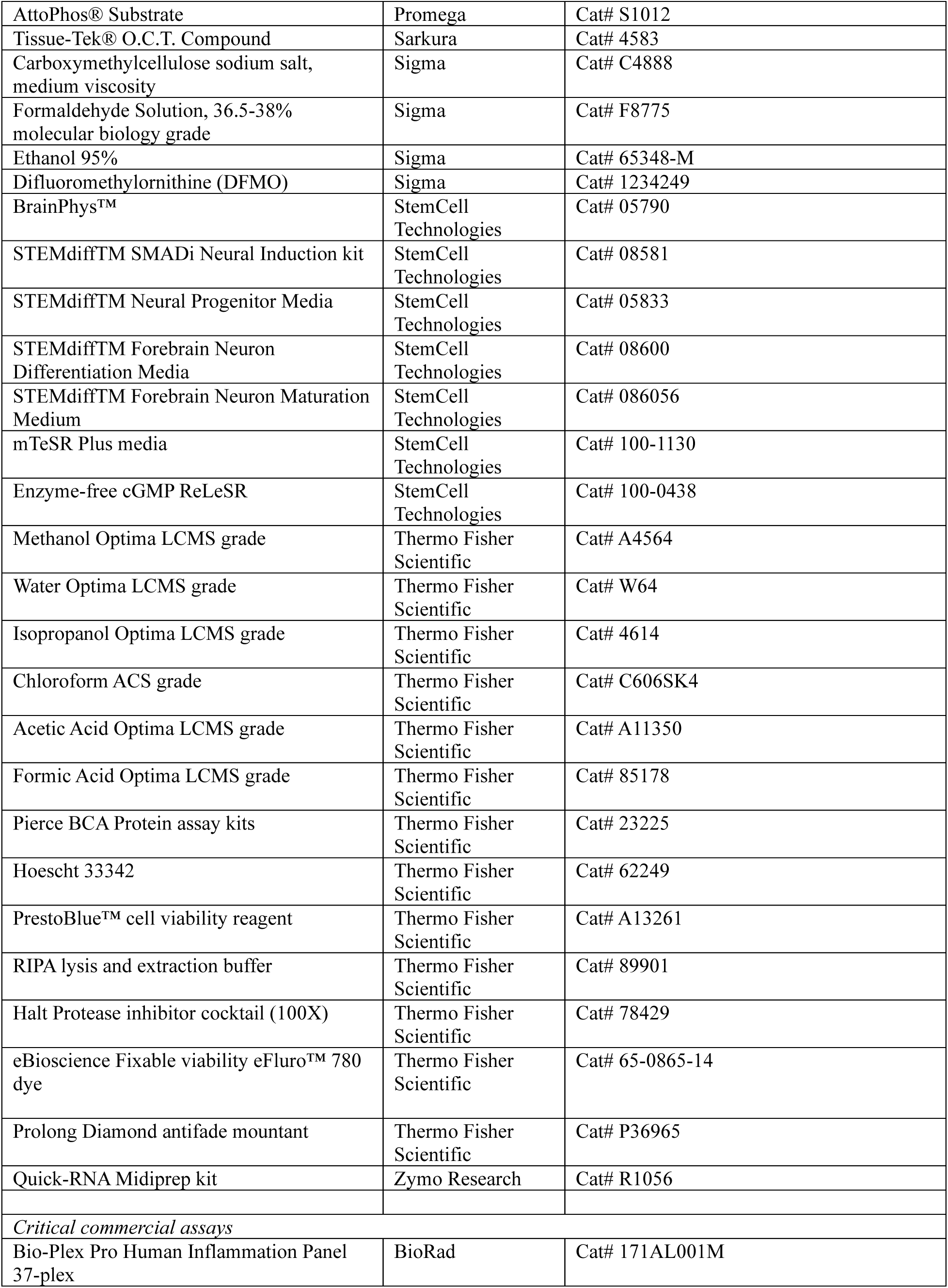
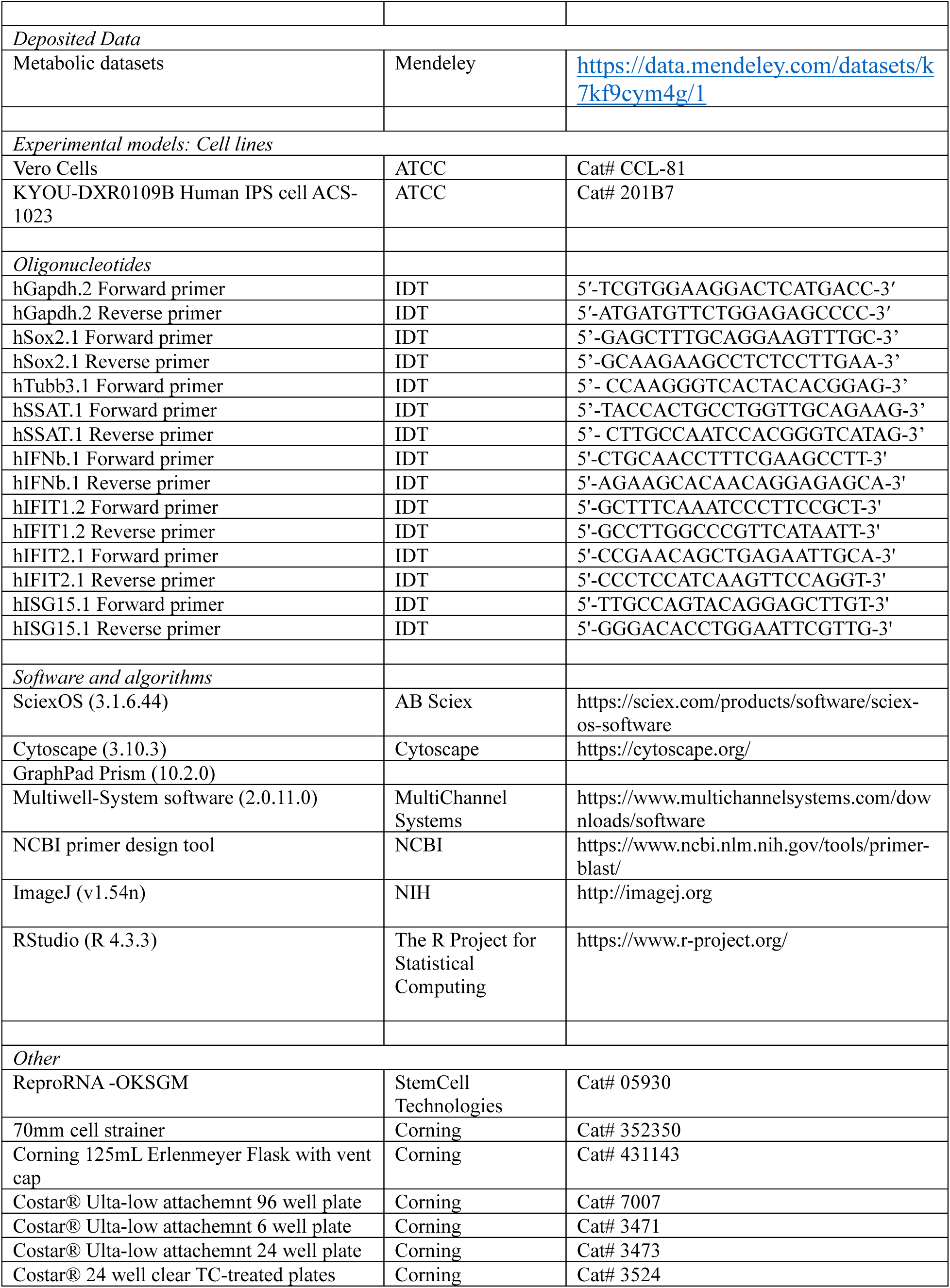
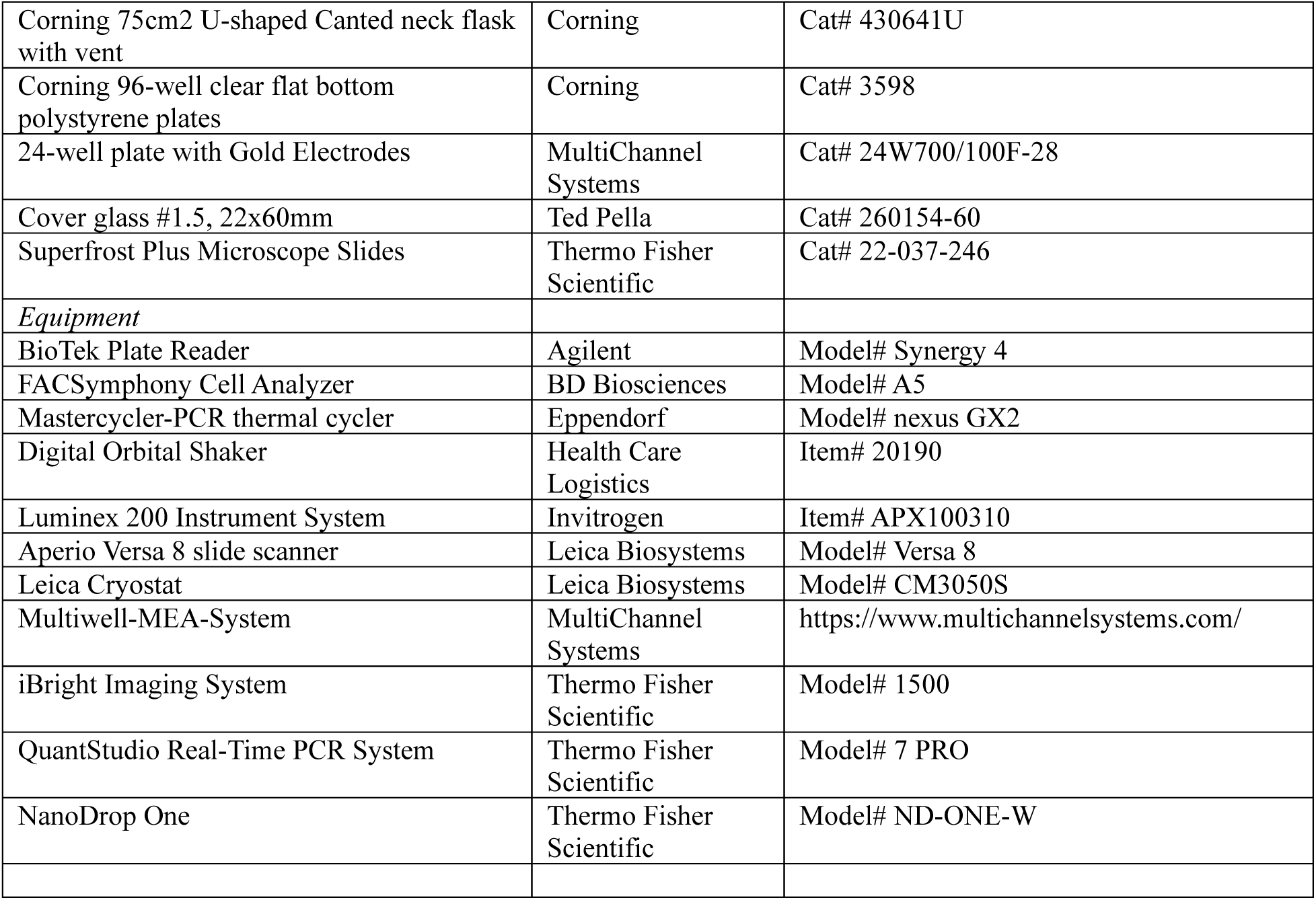

## Acknowledgments

This research was supported by the Intramural Research Program of the National Institutes of Health (NIH) through AI001102-11 (KEP) and AI001220-9 (RGM) The contributions of the NIH authors were made as part of their official duties as NIH federal employees, are in compliance with agency policy requirements, and are considered Works of the United States Government.

However, the findings and conclusions presented in this paper are those of the authors and do not necessarily reflect the views of the NIH or the U.S. Department of Health and Human Services. There was no input from funders in the study design, data collection and analysis, decision to publish, or preparation of the manuscript. We thank Sue Priola, Iain Fraser and Rachel Feldman for helpful review and discussion of the manuscript, Kader Cetin Gedik and Gina Montealegre for the patient care and data collection and Katherine Myint-Hpu for her support performing lumbar punctures on the patients.

## Data availability

Dataset information for all metabolomic studies including organoids, CSF and cells are available at: https://data.mendeley.com/datasets/k7kf9cym4g/1. Other data generated including real-time PCR analysis, protein expression and IHC are either included in this published article and supplemental figures or are available from the corresponding author upon request.

