## supplemental Tables and files for "Proteasome mutations associated with CANDLE syndrome cause altered neuronal development by dysregulating polyamine synthesis"

Table S1. Derivation of iPSCs used for organoid generation.

| *ID*[a] | *Diagnosis*[b] | *Mutation(s)*[c] | *iPSC source*[d] | *Source Ref*[e] | *Reprog*[f] | *Add Info*[g] | *Age*[h] | *Sex* | *symb*[i] |
|---|---|---|---|---|---|---|---|---|---|
| *CAN1* | CANDLE | PSMB8(T75M)/ PSMA3 digenic | patient skin fibroblast | | Episomal Vectors | ADHD | 11.8 | M | □ |
| *CAN2* | CANDLE | PSMB8 (T75M) | patient skin fibroblast | | Episomal Vectors | - | 12.5 | F | ○ |
| *CAN3* | CANDLE | PSMB8 (T75M/ T92A) | patient skin fibroblast | | Sendai virus | - | 24.8 | M | ◇ |
| *HC1* | Healthy | - | skin fibroblast | ATCC (1023) | Retroviral | - | 36 | F | ● |
| *HC2* | Healthy | - | skin fibroblast | RAH019A | Plasmid | - | 67 | F | ■ |
| *HC3* | Healthy | - | skin fibroblast | ASE-9209 | Episomal | - | 47 | F | ◆ |

a: identification of patient for this study

b: diagnosis of patient.

c: mutation associated with CANDLE

d: source of cells for generation of iPSCs

e: source reference for control iPSC lines

f: how cells were reprogrammed for iPSCs

g: additional information of patients. ADHD: Attention-deficit/hyperactivity disorder; NK, normal karyotype

h: Age at which iPSCs were generated

i: symbol used to reference this patient/iPSC/CO throughout the manuscript

Table S2. Age and Sex information for CSF samples

| ID | AGE | SEX | GENOTYPE |
|---|---|---|---|
| **HC3** | 29 | M | |
| **HC4** | 50 | M | |
| **HC5** | 43 | F | |
| **HC6** | 54 | F | |
| **HC7** | 50 | F | |
| **HC8** | 46 | M | |
| **HC9** | 41 | F | |
| **CAN3** | 20 | M | compound heterozygous *PSMB8* |
| **CAN6** | 3.5 | M | digenic *PSMB4, PSMB9* |
| **CAN7** | 7.5 | F | *PSMB8* dominant negative |
| **MS1** | 27 | F | |
| **MS2** | 74 | M | |
| **MS3** | 34 | F | |
| **MS4** | 47 | F | |
| **MS5** | 58 | F | |
| **MS6** | 58 | F | |
| **MS7** | 28 | M | |
| **MS8** | 59 | M | |
| **MS9** | 49 | M | |
| **MS10** | 67 | M | |

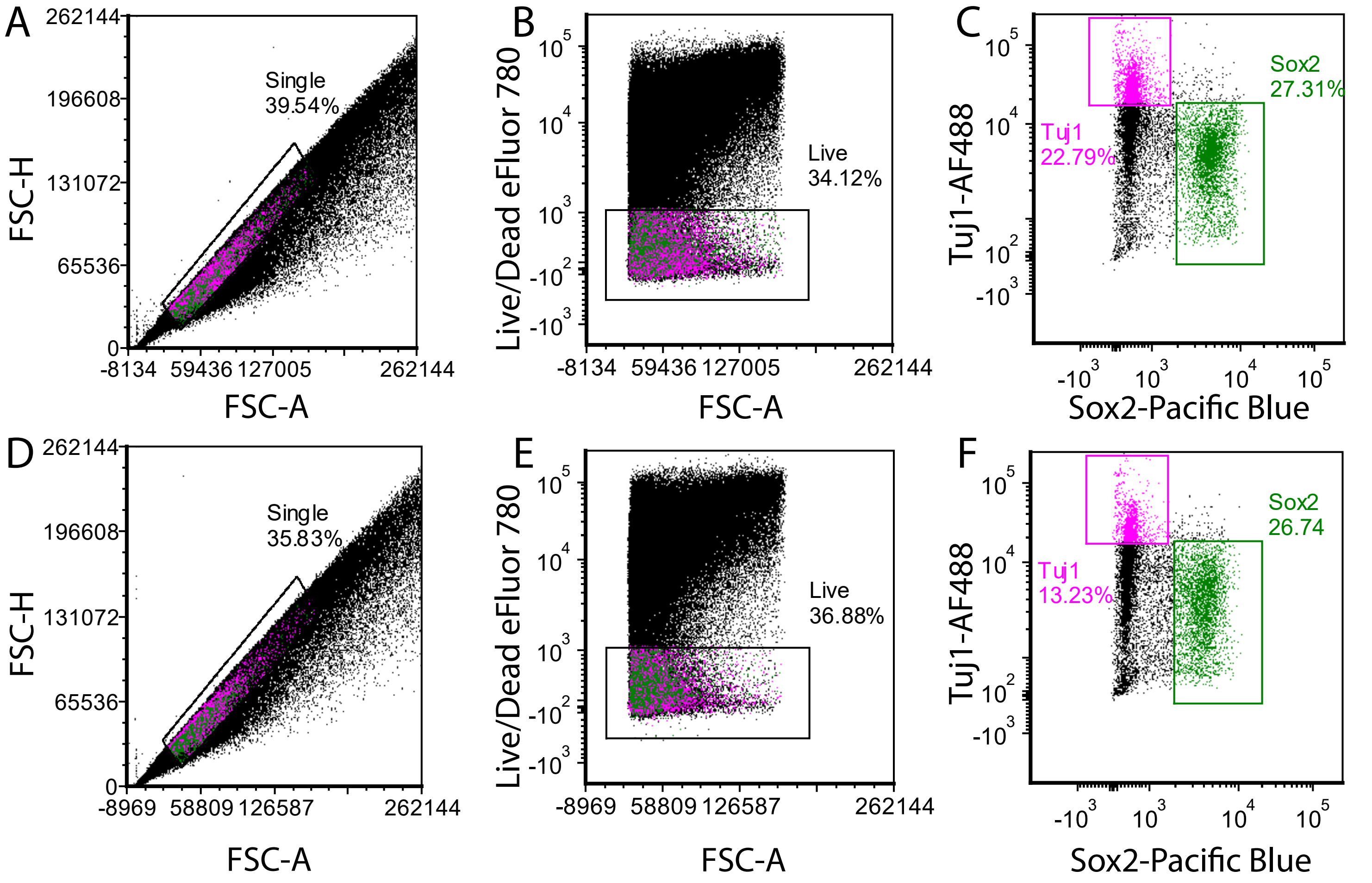


**Supplemental Figure 1. Flow cytometric gating strategy and analysis of cerebral organoids.** Flow cytometry was performed on n=6 dissociated COs from HC1 (A-C) and CAN1 (D-F). Doubles were removed from analysis by including events outside the diagonal when plotting FSC height x area (A&D). Live cells were identified by excluding labeling events labeled with Live/Dead eFluor 780 (B, E). Within the live cell population, Sox2 (green) and Tuj1 (magenta) immunolabeling was quantified (C&F).

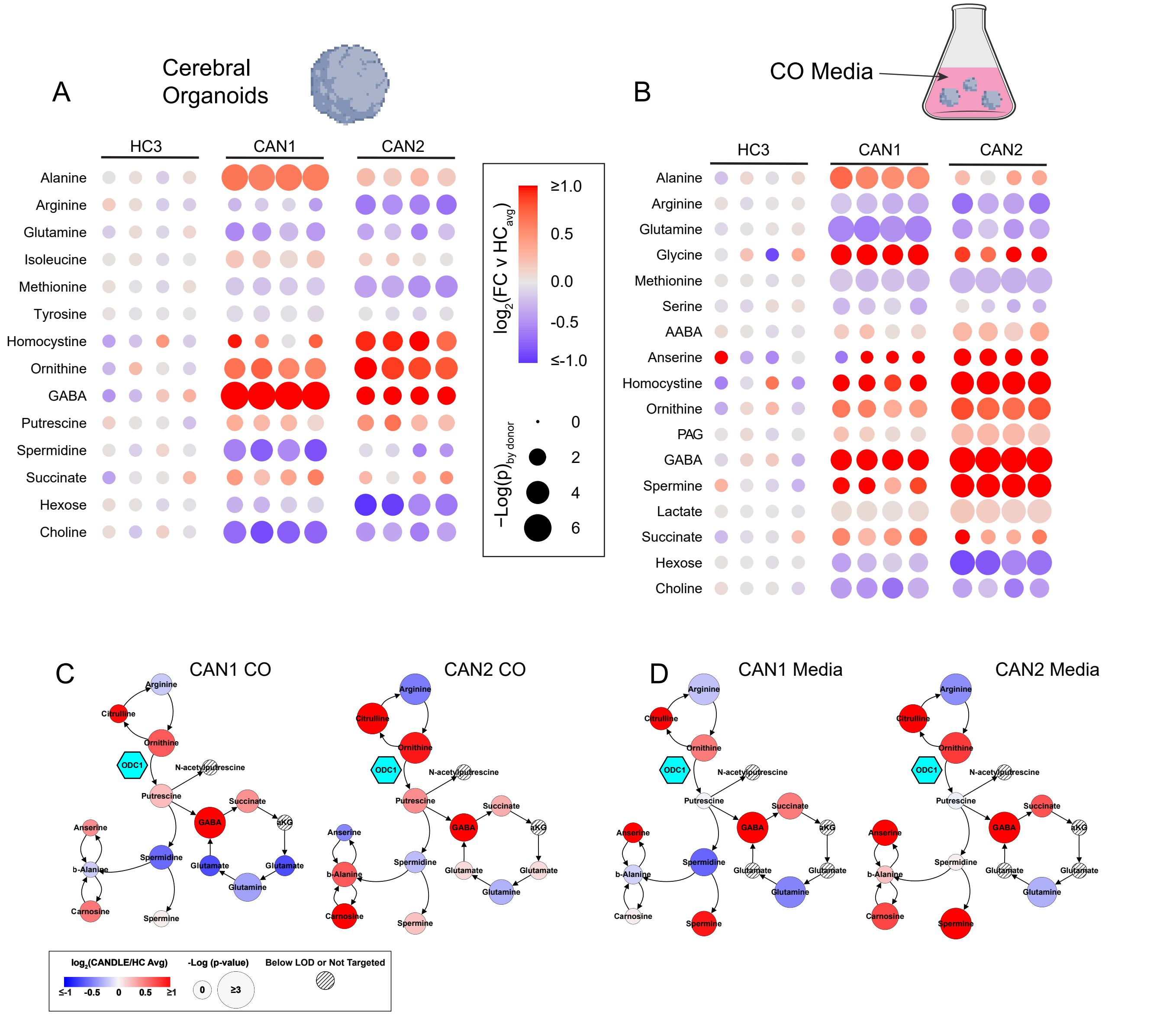


**Supplemental Figure 2. Biocrates metabolomics for COs and CO media.** A Biocrates Quant500 kit was used to measure metabolites from (A) COs and (B) CO media from HC3, CAN1, and CAN2. Displayed metabolites significantly varied between HC and CAN, with CAN1 and CAN2 pooled. Significance defined as passage of a 10 % FDR correction for multiple comparisons. In A&B the color of each node reflects the individual fold change (as the base two logarithm) for each CO analyzed compared to the average value of the HC3 COs for that metabolite. The size of each node reflects the negative logarithm of the raw p-value from a comparison of that patient to HC3 by a t-test. Only non-lipid metabolites were included in this analysis. (C&D) Amine metabolism maps of the same CO and CO-media metabolite comparisons displayed in (A&B) with the fold change vs. HC3 represented as color and the significance represented as node size as indicated in the legend. Hatch marked nodes indicate metabolites below the limit of detection or not targeted via this assay.

Supp Fig 3: Nucleobase and Acetyl-CoA metabolism

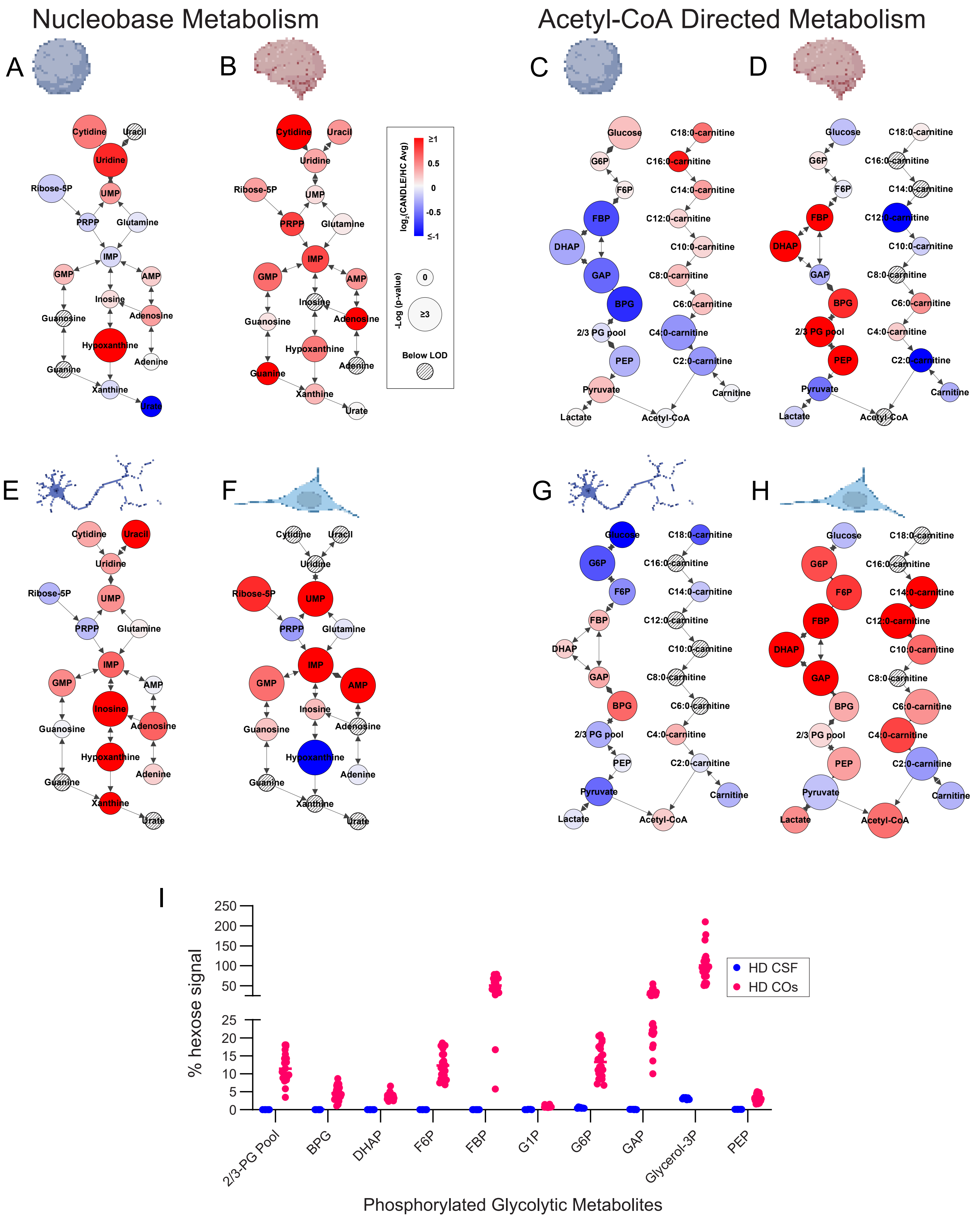

**Supplemental Figure 3. Cross matrix, CANDLE-driven nucleobase and acetyl-CoA directed metabolic patterns. (A&B)** Metabolic maps of CANDLE-associated changes in metabolite profiles compared to HCs in nucleobase metabolism and nucleobase salvage from **(A)** COs and **(B)** CSF. **(C&D)** Metabolic maps of CANDLE-associated changes in metabolite profiles compared to HCs in Acetyl-CoA generating pathways from **(C)** COs and **(D)** CSF. **(E&F)** Metabolic maps of CANDLE-associated changes in metabolite profiles compared to HCs in nucleobase metabolism and nucleobase salvage from **(E)** iPSC-derived neurons and **(F)** iPSC-derived NPCs. **(G&H)** Metabolic maps of CANDLE-associated changes in metabolite profiles compared to HCs in Acetyl-CoA generating pathways from **(G)** iPSC-derived neurons and **(H)** iPSC-derived NPCs. Across all maps, fold change (as the base two logarithm) of metabolites in CANDLE samples compared the HC samples is represented as color and the raw p-value (as the negative logarithm) via a t-test between CANDLE and HC is represented as node size. Replicates by sample type were: COs technical n=12 per donor, biological n=3 CAN n=2 HC, CSF biological n=3 CAN n=7 HC, iPSC-derived neurons and NPCs technical n=6, biological n=3 CAN n=2 HC. **(I)** Comparison of signal-levels by LC-MS/MS of phosphorylated glycolytic intermediates by matrix in HC samples. All displayed metabolite signals were normalized to the signal for hexose as a related non-energized metabolite.

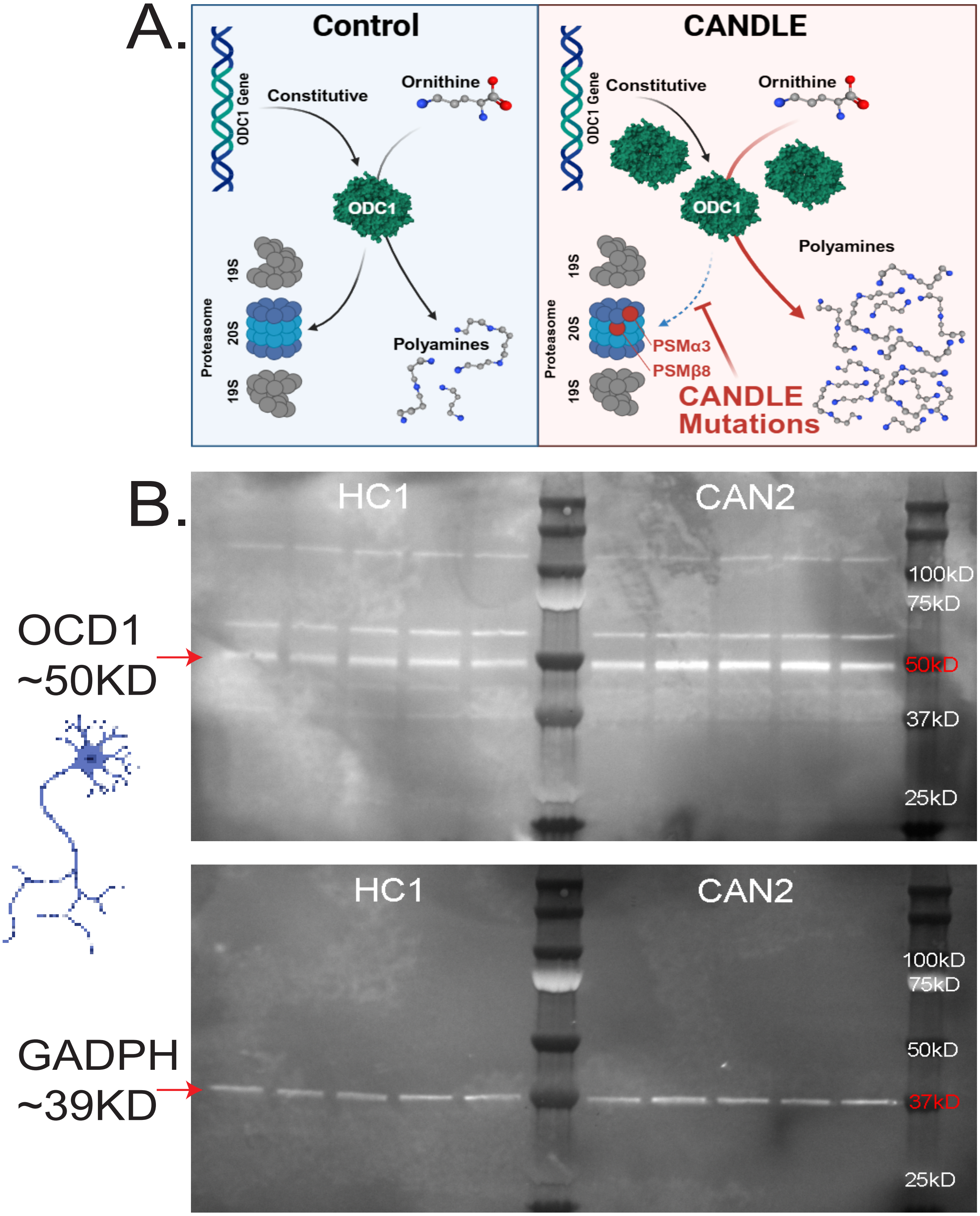


**Supplemental Figure 4. Whole SDS page gel of ODC1 expression in neurons.** (A) Graphical representation of the function and life cycle of ODC1 in normal (left) and CANDLE (right) cells. Under normal conditions, ODC1 is constitutively transcriptionally expressed and then degraded via the 20S subunit of the proteasome to regulate polyamine expression. CANDLE mutations impair this degradation leading to ODC1 and polyamine overexpression. (B) Antibody labeling for ODC1 protein in HC1 and CAN2 iPSC-derived neurons in the complete blot (top) with the primary band at ~59KD indicated in red. The same blot was reporbed for GAPDH expression (bottom) with the primary band at ~39KD.

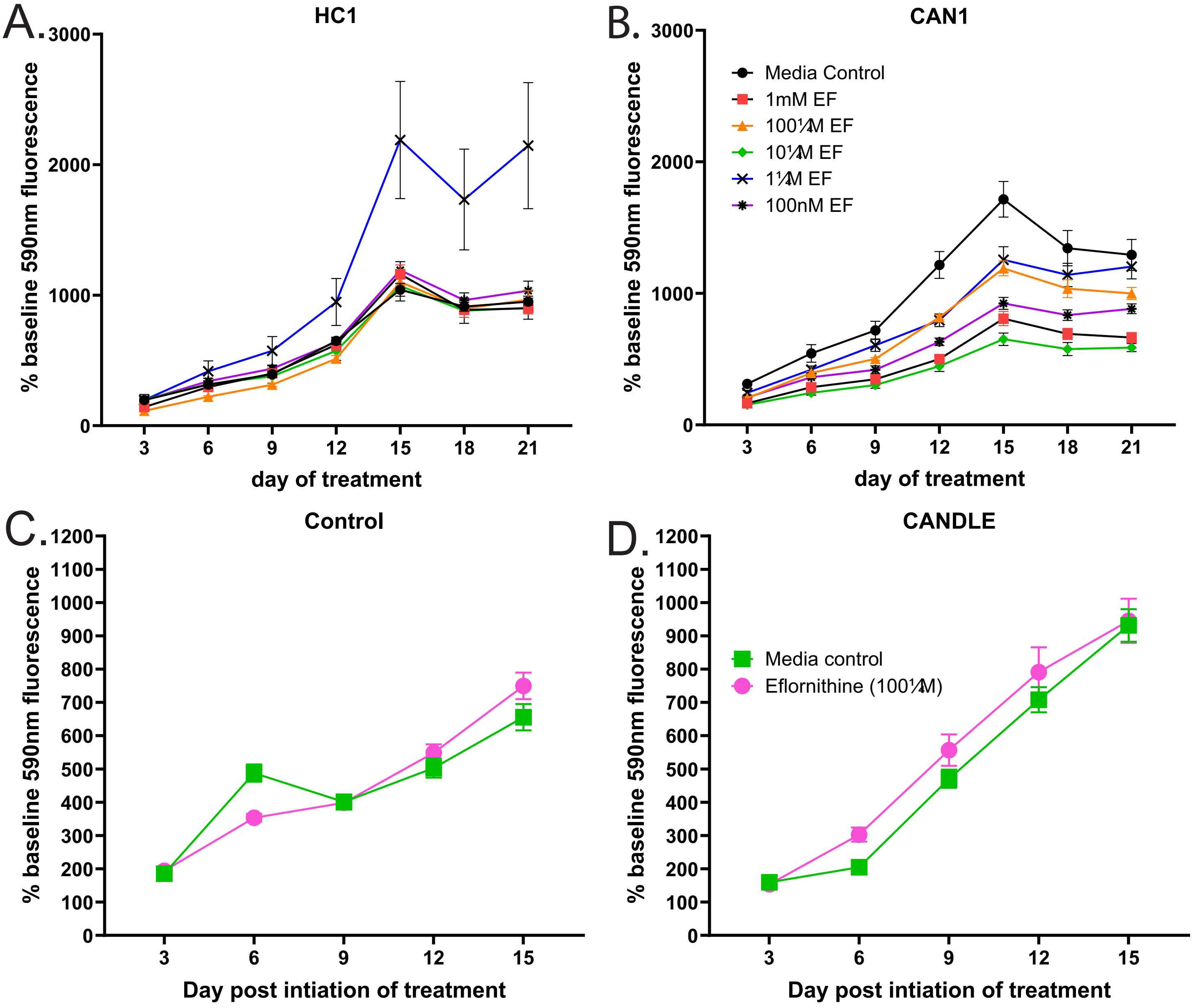


**Supplemental Figure 5. Toxicity assessment of Eflornithine in control and CANDLE organoids.** COs generated from n=3 HC1 (A) and n=3 CAN1 (B) were administered daily varying concentrations of DFMO in the culture media to determine the drug's effect on cellular reductive capacity as an indicator of cell health. The highest DFMO concentration with a similar level in cellular reductive capacity to media control (100mM) was selected for further metabolic and transcriptional experiments. Metabolic capacity of n=16 COs generated from HC1 and HC2 (D) and n=24 COs (E) generated from CAN1-3 used in metabolic and transcriptional experiments (Figure 7) throughout treatment.

Supplemental Fig 6

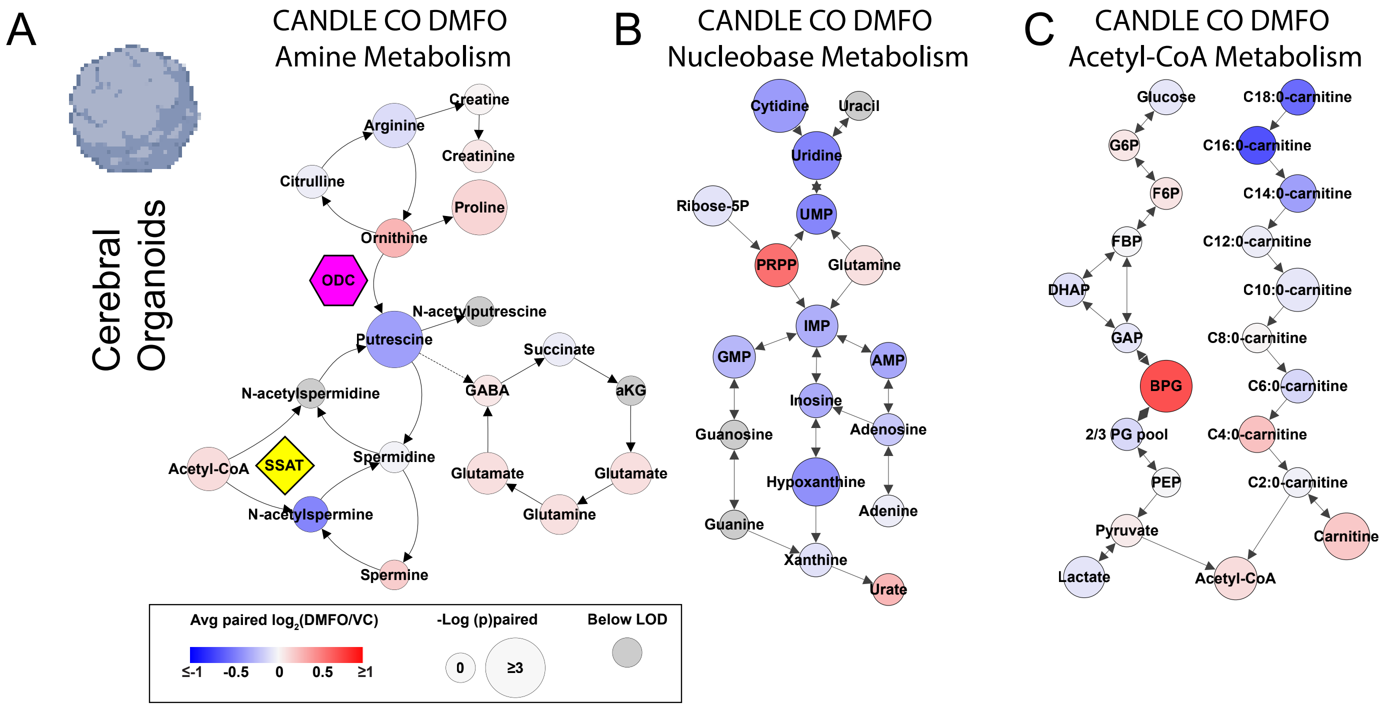


**Supplemental Figure 6. DFMO reverses CANDLE-associated amine, nucleobase, and Acetyl-CoA metabolic pattern in COs. (A)** A map of changes in amine metabolism associated with the paired comparison of donor metabolite averages for 100 µM DMFO-treated CANDLE COs vs. vehicle controls with the fold change of metabolites represented as color and the significance represented as node size (CO technical n=12, biological n=3 CAN). **(B)** A map of DMFO driven changes in CANDLE COs in nucleobase metabolism using the criteria described in **(A)**. **(C)** A map of DMFO driven changes in CANDLE COs in Acetyl-CoA generating pathways using the criteria described in A
